# Premalignant *Nf1, Trp53*-null Oligodendrocyte Precursor Cells Become Stalled in a Heterogeneous State of Replication Stress Before Gliomagenesis

**DOI:** 10.1101/2020.03.30.017228

**Authors:** Matthew D. Sutcliffe, Rui P. Galvao, Lixin Wang, Jungeun Kim, Lauren K. Rosenfeld, Shambhavi Singh, Hui Zong, Kevin A. Janes

## Abstract

Cancer evolves from premalignant clones that accumulate mutations and adopt unusual cell states to achieve transformation. Previously, we pinpointed the oligodendrocyte precursor cell (OPC) as a cell-of-origin for glioma, but the early changes of mutant OPCs during premalignancy remained unknown. Using mice engineered for inducible *Nf1–Trp53* loss in OPCs, we acutely isolated labeled mutant OPCs by laser-capture microdissection and determined gene-expression changes by bulk RNA sequencing and a fluctuation analysis, called stochastic profiling, which uses RNA-sequencing measurements from random pools of 10 mutant cells. At 12 days after *Nf1–Trp53* deletion, while bulk differences were mostly limited to mitotic hallmarks and genes for ribosome biosynthesis, stochastic profiling revealed a spectrum of stem-progenitor (*Axl, Aldh1a1*), proneural, and mesenchymal states as potential starting points for gliomagenesis. At 90 days, bulk sequencing detected very few differentially expressed transcripts, whereas stochastic profiling revealed cell states for neurons and mural cells that do not give rise to glial tumors, suggesting cellular dead-ends for gliomagenesis. Importantly, we identified mutant OPCs that strongly expressed key effectors of nonsense-mediated decay (*Upf3b*) and homology-dependent DNA repair (*Rad51c, Slx1b, Ercc4*) along with DNA-damage markers suggesting transcription-associated replication stress. Analysis of 10-cell transcriptomes at 90 days identified a locus of elevated gene expression containing an additional repair endonuclease (*Mus81*) and *Rin1,* a Ras–Raf antagonist and possible counterbalance to *Nf1* loss. At 150 days, *Rin1* was microdeleted in some gliomas and downregulated in all others. Replication stress may pose a considerable bottleneck that must be resolved for gliomas to initiate.

**Statement of significance:** In situ stochastic profiling of heterogeneous cell states in a mouse model of glioma uncovers regulatory confusion in a glioma cell-of-origin and defines a state of replication stress that precedes tumor initiation.

## Introduction

Glioblastoma multiforme (GBM) is the most common type of malignant brain tumor and remains incurable despite decades of basic and clinical research. There are multiple subtypes of GBM that differ in prevalence, mutation frequency, response to therapy, and cell-state composition (1). The latest conceptual frameworks support three subtypes, two of which encapsulate the proneural vs. mesenchymal regulatory state of GBM tumor cells (2). The third (“classical”) describes mesenchymal tumor cells mixed with astrocytes instead of the microglial recruitment characteristic of mesenchymal GBMs (3). Malignant cells of different subtypes often reside in the same GBM (4), and advanced GBM clones can give rise to cells with OPC-, neural progenitor-, mesenchymal-, and astrocyte-like signatures (5), suggesting that subtypes could reflect dynamic cell states in glioma. Interestingly, malignant cell-state trajectories appear to be simpler in lower-grade gliomas, even though these tumors also show a mixture of cell-state signatures (6). For multiple glioma subtypes, tumor ecosystems are maintained by a progenitor-like subpopulation that yields terminally differentiated progeny with oligodendrocytic or astrocytic features (7–9). It is unclear whether cell state heterogeneity is even simpler or more chaotic during premalignancy, as glioma-predisposed cells occupy various states until reaching a productive one for long-term tumor growth (10).

The Cancer Genome Atlas project identified three frequently mutated pathways in GBM: Arf/p53 signaling, receptor tyrosine kinase signaling, and the G1 checkpoint (11). Both patient sample analysis and genetically engineered mouse models (GEMMs) indicate oligodendrocyte precursor cells (OPCs) as a cell-of-origin for glioma (12–14). Toward modeling the very earliest steps of gliomagenesis, we previously developed a GEMM that incorporates OPC-selective conditional deletion of *Nf1–Trp53* together with lineage tracing to mark cells upon tumor-suppressor loss (15,16). Deletion in adult mice reactivates OPCs and restores proliferation to perinatal levels within 12 days. However, even though mutant OPCs show a net decrease in oligodendrocyte differentiation at 90 days after gene inactivation, proliferation rates decline to basal levels, right up until the onset of explosive gliomas. Besides these time-dependent changes in proliferation and differentiation, the cell-state changes during premalignancy remain unknown.

Here, we deconstructed the glioma GEMM’s premalignant phases by a combination of bulk transcriptomics and targeted analysis of heterogeneous cell states through 10-cell RNA sequencing (10cRNA-seq) (17–19). Both approaches used laser-capture microdissection (LCM) to isolate labeled cells in situ to avoid tissue dissociation-induced changes of gene expression would confound our analysis (20). Shortly after deletion–labeling, bulk transcriptomic differences largely indicated generic adaptations to *Nf1–Trp53* loss, whereas the heterogeneity analysis of transcriptional variation gave clues about the early states of labeled OPCs. Interestingly, intermediate premalignancies showed very limited differences in the bulk analysis, but stochastic profiling by 10cRNA-seq revealed substantial increases in heterogeneity of cell states indicated by terminal markers of aberrant differentiation. Among these observations, we discovered a collection of cells that was not deconvolvable into any recognized cell types of the brain. These samples marked a cell state characterized by elevated DNA repair and nonsense-mediated decay, suggesting difficulty in resolving the transcription-associated replication stress caused by *Nf1–Trp53* deletion. Our results support a conceptual model of early gliomagenesis that is chaotic and largely unproductive. Even with two glioma-predisposing deletions in two tumor-suppressor genes, *Nf1,p53-null* mutant OPCs often end up in aberrant “dead-ends” for tumorigenesis or frozen in a state of replication stress. Glioma precursors appear to struggle in bypassing proliferation–differentiation roadblocks, which might need to be surmounted concurrently through extensive homology-independent rearrangements (21) that enable progression into full-blown tumors.

## Materials and Methods

### Tissue sources

All mice were maintained according to practices prescribed by the National Institutes of Health in accordance with the IACUC protocol #3955. All animal procedures were approved by the Animal Care and Use Committee at the University of Virginia, accredited by the Association for the Assessment and Accreditation of Laboratory Animal Care (AAALAC). The following mouse strains were crossed on a mixed background: Cspg4 (Ng2)-CreER (a kind gift from Dr. Akiko Nishiyama), p53^KO^ (stock #002101, Jackson Laboratories), Rosa-tdTomato (stock #007908, Jackson Laboratories), NF1^flox^ (strain #01XM4, National Cancer Institute), p53^flox^ (strain #01XC2, National Cancer Institute). Genotypes of the mice used in the bulk analysis were *Cspg4-CreER; Rosa-tdTomato; Nf1^flox/flox^; p53^-/flox^* (mutant) and *Cspg4-CreER; Rosa-tdTomato* (control). Genotypes of the mutant mice used in the 10cRNA-seq analysis were *Cspg4-CreER; Rosa-tdTomato; Nf1^flox/flox^; p53^flox/flox^*. Genotyping was performed by PCR as described (15). Tamoxifen citrate tablets (10 mg/tablet, Mylan) were crushed, resuspended in water, and delivered by oral gavage. All mutant mice were given daily doses of 200 mg tamoxifen/kg for five consecutive days starting at P45. Control mice were given tamoxifen at the end of each experiment instead of at P45 to ensure labeling of OPCs rather than mature oligodendrocytes.

Mouse brain tissue was isolated at either 12, 90, or 150 days post-induction (dpi) from the start of tamoxifen treatment. Animals were not randomized by time point but balanced to the extent possible by sex. For the bulk samples, isopentane was pre-chilled for ~30 min on dry ice to −50°C or lower. To dissect brains, mice were transcardially perfused with PBS supplemented with Procaine and Heparin (Sigma-Aldrich), then brains were dissected and slowly submerged in the pre-chilled isopentane. Brains were removed after one minute, wrapped in Parafilm and stored at −80°C until ready for sectioning and LCM. For the 10cRNA-seq samples, the freshly dissected brain hemispheres were separated and the olfactory bulbs were isolated. Each olfactory bulb was freshly cryoembedded in a dry ice-isopentane bath and stored at −80°C wrapped in aluminum foil until sectioning and LCM.

Before cryosectioning the bulk samples, untreated, plain glass slides were irradiated for 30 minutes with an ultraviolet lamp to eliminate RNases and the cryostat was cleaned with ethanol and RNase-Away (Sigma-Aldrich). Cryosections were 10 μm thick. Once a section was collected, the slide was kept inside the cryostat to avoid thawing and water condensation. For the 10cRNA-seq samples, samples were equilibrated to −24°C in a cryostat before sectioning. 8-μm sections were cut and wicked onto Superfrost Plus slides. To preserve tissue integrity, slides were pre-cooled on the cutting platform for 15–30 sec before wicking, and the section was carefully placed atop the cooled slide. Then, the slide was gently warmed from underneath by tapping with a finger until the section was minimally wicked onto the slide. All wicked slides were stored in the cryostat before transfer to −80°C storage on dry ice. Frost build-up was minimized by storing cryosections in five-slide mailers.

### Fluorescence-guided LCM

For the bulk samples, frozen slides were immediately placed in pre-chilled (−20°C) 100% ethanol for five minutes, then transferred to room temperature 100% xylene for five minutes twice. Finally, slides were removed from xylene and air-dried for five minutes. For the 10cRNA-seq samples, fluorescence-guided LCM was performed as described previously (18). Immediately from −80°C storage, slides were fixed in 70% ethanol for 15 seconds, 95% ethanol for 15 seconds, and then 100% ethanol for two minutes before clearing with xylene for two minutes. Slides were air-dried for 5–10 minutes before beginning LCM. Freshly fixed samples were microdissected on an Arcturus XT LCM instrument (Applied Biosystems) using Capsure HS caps (Arcturus). The smallest spot size on the instrument (10–12 μm) captured one OPC per laser shot.

### RNA extraction and amplification

For bulk samples, total RNA from the Capsure HS cap was extracted and purified with the PicoPure RNA isolation kit (Arcturus) according to manufacturer instructions. Briefly, tissue was immediately incubated in 10 μl extraction buffer at 42°C and stored at −80°C. Samples were column purified and digested with DNase to remove genomic DNA. Total RNA was eluted in 11 μl RNase-free water and concentrated to 5 μl in a speedvac for five minutes at room temperature. Single-primer, isothermal linear amplification (Ribo-SPIA) of purified total RNA was performed with the Ovation RNAseq System V2 kit (NuGen) according to manufacturer instructions. Amplified cDNA molecules were purified before generating sequencing libraries.

For 10-cell samples, RNA extraction and amplification of microdissected samples was performed as described previously (18). Briefly, RNA was eluted from Capsure HS caps (Arcturus) by digestion of microdissected cells with proteinase K. Biotinylated-cDNA was then synthesized from RNA eluted from captured cells and purified using streptavidin magnetic beads (Pierce) with a 96 S Super Magnet Plate (Alpaqua). Residual RNA was degraded by RNAse H (NEB) digestion and poly(A) tailed with terminal transferase (Roche). Poly(A)-cDNA was then amplified using an AL1 primer (ATTGGATCCAGGCCGCTCTGGACAAAATATGAATTCTTTTTTTTTTTTTTTTTTTTTTTT) and a blend of Taq polymerase (NEB) and Phusion (NEB).

### Quantitative PCR (qPCR)-based quality assessment

Detection of transcripts by qPCR was performed on a CFX96 real-time PCR instrument (Bio-Rad) as described previously (22). Samples were filtered based on three loading controls: *Rplp1, Gapdh,* and *Apoe.* Samples were retained if the geometric mean of the *Rplp1–Gapdh– Apoe* quantification cycles was within 3.5x interquartile range of the median; samples outside that range were excluded because of over- or under-capture during LCM. We also sought to exclude samples that were overly contaminated with *Cspg4*-expressing pericytes. Therefore, samples with detectable *Pdgfrb, Des,* and *Anpep* were excluded from sequencing. qPCR primer sequences are available in Supplementary Table ST1.

### Library preparation and sequencing

For bulk samples, Illumina sequencing libraries were made by standard methods; briefly, double-stranded cDNA was mechanically sheared to 150–600 bp by sonication using a Bioruptor (Diagenode), and the fragments end-repaired, A-tailed and ligated to T-overhang adaptors. Samples were ligated to Illumina adapters each with a 5-nt index, amplified for 10 to 12 cycles, and size selected to between 250–500 bp. Samples were sequenced on an Illumina HiSeq 2000 instrument.

10cRNA-seq libraries for sequencing were re-amplified, purified, and tagmented as described previously (18). Briefly, each poly(A) PCR cDNA sample was re-amplified for a number of cycles where the amplification remained in the exponential phase (typically 10 to 20). Re-amplified cDNA was then twice purified with Ampure Agencourt XP SPRI beads. After bead purification, samples were quantified on a CFX96 real-time PCR instrument (Bio-Rad) using a Qubit BR Assay Kit (Thermo Fisher). Samples were diluted to 0.2 ng/μl and tagmented with the Nextera XT DNA Library Preparation Kit (Illumina). Samples were sequenced as previously described (18). Samples were multiplexed at an equimolar ratio, and 1.3 pM of the multiplexed pool was sequenced on a NextSeq 500 instrument with NextSeq 500/550 Mid/High Output v1/v2/v2.5 kits (Illumina) to obtain 75-bp paired-end reads.

### RNA sequencing alignments

Adapters were trimmed using fastq-mcf (version 1.1.2-779) in the EAutils package with the following options: -q 10 -t 0.01 -k 0 (quality threshold 10, 0.01% occurrence frequency, no nucleotide skew causing cycle removal). Quality checks were performed using FastQC (version 0.11.8) and MultiQC (version 1.7). Datasets were aligned to the mouse transcriptome (GRCm38.82) along with reference sequences for ERCC spike-ins, using RSEM (version 1.3.0) and Bowtie 2 (version 2.3.4.3). RSEM options for the 10cRNA-seq data also included the following options: --single-cell-prior --paired-end. RSEM read counts were converted to transcripts per million (TPM) by dividing each value by the total read count for each sample and multiplying by 10^6^. For bulk samples with multiple library preparations, RSEM read counts were combined before TPM calculation. Mitochondrial genes and ERCC spike-ins were not counted towards the total read count during TPM normalization.

### Differential expression analysis

DESeq2 (version 1.26.0) was used to identify differentially expressed transcripts. For the bulk samples, outliers were first identified by principal components analysis and removed from subsequent analysis. The differential expression analysis was performed separately at each of the three timepoints. Mouse sex was included as a covariate with full model design “~ genotype + sex” and reduced model design “~ sex”. Genes with an adjusted *p* < 0.05 were considered significantly up- or downregulated between the control and mutant samples. For the 10cRNA-seq samples, DESeq2 was used to identify differentially expressed transcripts between the U group and the O–E–N union group with model design “~ group”. Genes with an adjusted *p* < 0.05 were considered significantly up- or downregulated between the two groups.

### Overdispersion-based stochastic profiling

Stochastic profiling with 10cRNA-seq data was performed separately for the male and the female datasets exactly as described in the accompanying contribution (19).

### Robust identification of transcriptional heterogeneities through crossvalidation

To identify heterogeneously expressed genes that were robust to sample outliers, we performed 100 leave-one-out simulations separately on the male and female datasets. For each simulation, one 10-cell sample and one pooled control were randomly chosen and removed from the dataset. To maintain a total of 28 10-cell samples and 20 pooled controls, one each of the remaining 10-cell samples and pooled controls was duplicated. The overdispersion analysis was performed as described above, returning a list of heterogeneously expressed genes for each simulation. Sex-specific heterogeneities were defined as those genes found heterogeneously expressed in at least 75% of the simulations.

### Glioma subtyping and single-cell inference

Bulk and 10cRNA-seq samples were projected onto the first two principal components derived from a C1-based GBM principal components analysis (3). Mouse genes were first converted into their human orthologs using the “biomaRt” package in R. Of the 714 human genes contributing to the principal components, 629 genes had mouse orthologs in our dataset that were used for the projection. TPM values were scaled to TPM/100 + 1 and log_2_-transformed before applying the projections. Maximum-likelihood inference using the “stochprofML” package in R was performed on the 10cRNA-seq 12 dpi sample scores in the first principal component to determine constituent single-cell contributions (23). Scores were right-shifted by 120 for parameterization of lognormal distributions; then, location parameters were left-shifted by 120 for display. The inference was performed with two lognormally-distributed cell populations and ten cells per sample. The probability density function for the mixed samples was determined by repeatedly sampling from the resulting single-cell distributions. To compare bulk samples and 10cRNA-seq samples along the proneural–mesenchymal and proliferation principal components at 12 dpi, 70–100 10-cell samples were randomly selected with replacement. TPM values were averaged before projecting onto the two principal components as described above.

### Lineage and cell-state deconvolutions

CIBERSORT was used to unmix cellular samples into their composite cell types (24). Cell-type signatures were derived using cell-sorted RNA-seq data (25). The two biological replicates from each of five cell types (astrocyte, endothelial, myelinating oligodendrocyte, microglia, and neuron) were averaged. Because of ambiguity in the oligodendrocyte-lineage samples and because differentiated OPCs would continue to be fluorescently labeled, one OPC sample and one newly-formed oligodendrocyte sample from (25) were averaged to yield a hybrid “OPC” more characteristic of the expected population of labeled cells. Signature genes were defined as the transcripts with the highest fold enrichment of one cell type vs. the average of the remaining cell types. Only genes with FPKM > 20 were included in the analysis. We identified 40 signature genes per cell type, and FPKM values were converted to TPM before the CIBERSORT analysis. To account for pericyte labeling in the cell unmixing analysis, the endothelial signature was exchanged for a similarly derived pericyte signature based on mural cell RNA-seq data (26). The new signature matrix was optimized as described above. CIBERSORT was run in relative mode or absolute mode where indicated.

### Immunofluorescence and image segmentation

Immediately from −80°C storage, frozen slides were fixed in prechilled (−20°C) 75% ethanol for 30 seconds and then post-fixed in 3.7% paraformaldehyde in PBS for 15 minutes at room temperature. Post-fixed slides were washed three times in PBS for five minutes each and then blocked with the basic M.O.M. immunodetection kit (Vector Laboratories) as recommended by the manufacturer. Primary antibody incubations were performed overnight at room temperature in M.O.M. diluent. Primary antibodies recognizing the following proteins were used: Ki67 (Thermo Fisher #MA5-14520, 1:500), phospho-Rb Ser^807/811^ (Cell Signaling #8516, 1:1600), Tgfbi (Novus #H00007045-B01P, 1:200), Mcl1 (Santa Cruz #sc-819, 1:200), Fn1 (BD Biosciences #610077, 1:200), Actn1 (Cell Signaling #3134, 1:50), Vcam1 (Santa Cruz #sc-8304, 1:200), Ercc4 (Santa Cruz #sc-136153, 1:20), Upf3b (Thermo Scientific #PA5-51652, 1:50), phospho-H2A.X Ser^139^ (Cell Signaling #9718, 1:400 or Millipore-Sigma #05-636, 1:200). After washing, slides were incubated with Alexa Fluor 488- or 647-conjugated secondary antibodies, washed, counterstained with DAPI, quenched for autofluorescence, and mounted as described previously (27). Images were collected on 15–30 fields containing tdTomato^+^ mutant cells for each animal, and 3–4 animals were used per time point.

Image segmentation and single-cell quantification were performed with CellProfiler 3.0 (28) as follows. DAPI-positive nuclei were smoothed with a Gaussian filter and segmented by global two-class thresholding by Otsu’s method. tdTomato^+^ cells were selected among segmented nuclei with a median smoothed tdTomato fluorescence intensity above a fixed threshold for a given image set. Nuclei were dilated by 10 pixels (3.2 μm) to capture the vicinity of each cell, and median object intensities were measured in the Alexa Fluor 488 and Alexa Fluor 647 channels for analysis. If necessary, object intensities were downscaled to match the background single-cell distribution of the image set with the lowest background median fluorescence. Median object intensities were aggregated in MATLAB and fit to a lognormal distribution or a mixture of lognormal distributions with a shared log variance. Median intensity gates for positive or high cells were set at the 99^th^ percentile of the rightmost distribution.

### Regional inference of transcriptional activity and copy-number variation

Locally elevated gene expression and copy-number changes were estimated by inferCNV (https://github.com/broadinstitute/inferCNV) as described in the accompanying contribution (19). The normal reference samples for 10cRNA-seq data were scRNA-seq profiles or 10-cell pools from GSE60361 (29) and GSE75330 (30). The normal reference samples for bulk RNA-seq were control RNA-seq data collected at 12 dpi, 90 dpi, and 150 dpi.

### UMAP projections

All UMAP projections were generated using the R package “uwot” (version 0.1.5). For the 12 dpi UMAP projections, RSEM values were converted to TPM values before log_2_ scaling. Default parameters were used. For the combined bulk + 10cRNA-seq UMAP projections, RSEM values were converted to TPM values before log_2_ scaling. Bulk samples and 10cRNA-seq samples were then z-scored independently. The following UMAP parameter was used: n_neighbors = 20.

### Statistics

Sample sizes for stochastic profiling were determined by Monte Carlo simulation (31). Animal sizes for the bulk analysis provided 66–93% power to detect an effect size of two standard deviations according to noncentral *t* statistics. The significance of marker-positive populations identified by immunofluorescence was assessed by the “binocdf” function in MATLAB with a background probability of 1% corresponding to the positive gate at the 99^th^ percentile. Differences in median immunofluorescence between groups were assessed by the “ranksum” function in MATLAB. Differences in genes detected per sample between 10cRNA-seq and scRNA-seq were assessed by Kolmogorov-Smirnov test using the “ks.test” function in R. Significance of overlap was assessed by Fisher’s exact test using the “fisher.test” function in R. In the male and female overlaps, the total number of genes was defined by the intersection of filtered genes in the male and female datasets (10,585 genes for 12 dpi and 11,281 genes for 90 dpi). In the RHEG and DE overlaps, the total number of genes was defined by the intersection of the previously filtered gene list and the detected genes from the DE analysis (10,092 genes for 12 dpi and 10,503 genes for 90 dpi). Significance of the relative size of RHEG or DE gene lists was assessed by binomial test using the “binom.test” function in R. Hierarchical clustering was performed using “pheatmap” in R with Euclidean distance and “ward.D2” linkage. Venn diagrams were visualized using the “matplotlib_venn” package in Python. Gene set enrichment analyses were performed through the Molecular Signatures Database (32). Overlaps between gene lists and hallmark gene sets were computed using a hypergeometric test with false-discovery rate correction for multiple comparisons.

### Data availability

Bulk and 10cRNA-seq data from this study is available through the NCBI Gene Expression Omnibus (GSE147360, https://www.ncbi.nlm.nih.gov/geo/query/acc.cgi?acc=GSE147360 Reviewer token: odgbkqqmlxgdrof).

## Results

### Comparing bulk-transcriptome and cell-state changes in a GEMM of glioma

We previously described a GEMM for low-grade glioma and secondary GBM in which the glioma-related tumor suppressors *Nf1* and *Trp53* are deleted with a tamoxifen-inducible Cre (CreER) (15) under the control of a *Cspg4* promoter (*P_Cspg4_*) that specifically targets OPCs (12). *Nf1* deletion mimics receptor tyrosine kinase signaling; *Trp53* deletion disrupts tumor suppression through the Arf/p53 pathway (11). A Cre reporter is incorporated into the GEMM so that mutant OPCs express tdTomato for fluorescence visualization. Serial administration of tamoxifen creates tdTomato^+^ OPCs with an *Nf1*^Δ/Δ^*Trp53*^Δ/Δ^ mutant genotype throughout the brain (**Fig. 1A**). However, only 1–3 clones per animal expand as malignant gliomas in ~150 days (15), leaving open the question of what hurdles these clones must overcome.

**Figure 1.**
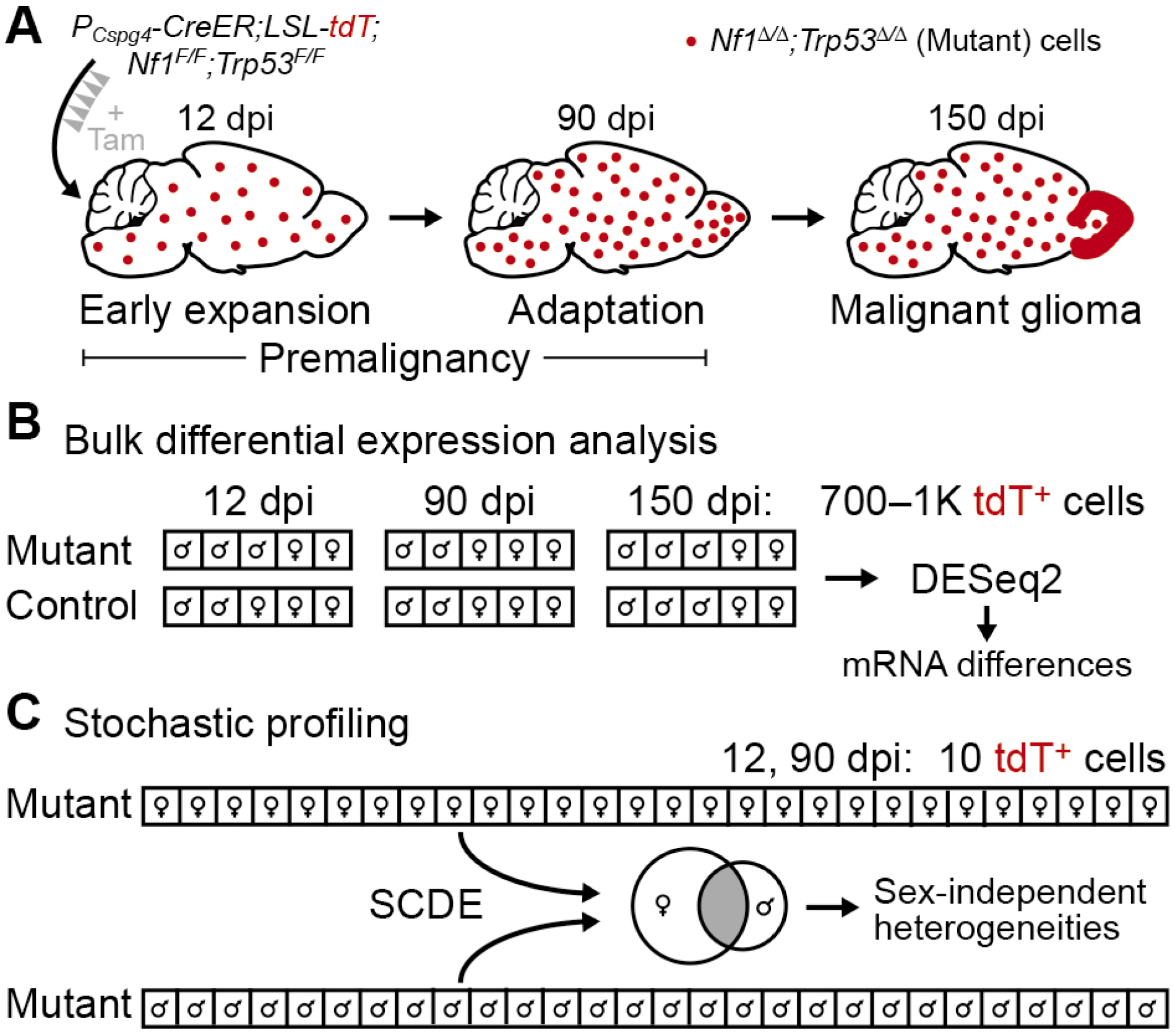
Transcriptomic analysis of bulk expression differences and regulatory heterogeneities in a GEMM of glioma. **A,** Summary of the glioma GEMM. Floxed alleles of *Nf1* and *Trp53* are inducibly deleted by tamoxifen (Tam) activation of CreER localized to oligodendrocyte precursor cells with a Cspg4 promoter (*P_Cspg4_*). The resulting *Nf1’’Trp53’’* (mutant) cells are labeled with tdTomato (tdT) and expand in the days post-induction (dpi) to form gliomas preferentially in the olfactory bulb (15). **B,** Experimental plan for bulk differential expression analysis. Large numbers of labeled cells from mutant and control animals were collected by LCM at the indicated dpi, linearly amplified, and sequenced for differential expression (*n* = 4–7 animals at roughly equal male-female proportion for each group). **C,** Experimental plan for heterogeneity analysis by stochastic profiling (17,19). Many 10-cell pools of mutant cells (*n* = 28 of each sex) were collected, measured by 10cRNA-seq (18), and assessed for regulatory heterogeneity with the abundance-dependent dispersion module of the SCDE package (42,43). Male and female candidates were intersected to arrive at the final set of regulatory heterogeneities.

To gain insight into the cell-state changes in mutant OPCs before gliomagenesis, we devised a study in which mutant OPCs were purified at two premalignant stages for geneexpression analysis: 12 dpi, when mutant OPCs show evidence of proliferative expansion (15); and 90 dpi, when mutant OPCs appear largely quiescent and are presumably adapting to prolonged loss of *Nf1* and *Trp53.* Given sex differences in the prevalence and molecular subtypes of human glioblastoma (33), we sought a balanced design that split observations evenly between males and females (**Fig. 1B** and **1C**). 150 dpi mutant samples containing frank gliomas were used to assess the fidelity of cell isolation and as a terminal reference point for the tumor population after transformation had occurred (**Fig. 1A** and **1B**).

Accessing labeled cells required some form of OPC isolation or purification with singlecell resolution. However, an earlier attempt to enrich OPCs by flow sorting (30) yielded fewer transcript alignments compared to other methods (18). Therefore, we used fluorescence-guided LCM under conditions of cryoembedding, sectioning, and fixation that retained tdTomato fluorescence and RNA integrity in labeled cells (18,34). We reasoned that *Nf1–Trp53* deletion would cause population-wide changes in gene expression compared to control OPCs as well as single-cell variations in transcript abundance requiring the stochastic-profiling method developed by our group (17,19). For the bulk analysis, we microdissected many hundreds of cells to minimize the impact of cell-to-cell variation (Supplementary Fig. S1) (31) and sequence without any exponential amplification that could skew transcript abundances (see Materials and Methods; **Fig. 1B**). The stochastic-profiling implementation, by contrast, involved repeated, random collection of 10-cell pools (17), which were preamplified, sequenced, and assessed for overdispersion as described in the accompanying contribution (19) (**Fig. 1C**). By applying different unbiased methods to the same microdissected subpopulation, our goal was to assess their relative merits for deciphering premalignancy.

### LCM-based RNA sequencing validates differential gene expression and proneural subtype of the glioma model

According to bulk microarray profiles and an early glioma-classification system (1), the glioma GEMM is most similar to the human proneural subtype (15). Subsequent work at the single-cell level suggested that intratumor heterogeneity of advanced gliomas is substantial (4), with the proportion of different cell types (recruited astrocytes vs. microglia) and cell states (proneural vs. mesenchymal) dictating the bulk classification. Glioma-propagating cells are now proposed to reside on a spectrum of proneural-to-mesenchymal cell fate, which is orthogonal to a cell’s proliferative state (3). The question of subtyping is also separate from differential expression, as many of the markers delineating a subtype will reflect vestiges of the cell type in which the tumor appeared. Together, these considerations motivated a reassessment of 150 dpi glioma cells by using the latest experimental and analytical methods (**Fig. 1B**).

Although gliomas were visibly macrodissectable at 150 dpi (**Fig. 2A**), mutant OPCs at earlier time points remained sparse throughout the tissue (**Fig. 2B**); thus, LCM was used for all sample types throughout the study. Several hundred laser shots of ~one-cell diameter could be fired per cryosection to generate enough material for RNA sequencing after Ribo-SPIA linear amplification (see Materials and Methods). We confirmed that transcriptomes of separate LCM bulk samples from the same tumor were highly correlated after linear amplification (Supplementary Fig. S2). Control OPCs were singular and much more entangled with unlabeled neighbors compared to prior applications of fluorescence-guided LCM by our group (20,34). To assess overall purity of normal controls, we used a collection of brain cell type-enriched marker transcripts (25) to construct a signature matrix for bulk sample deconvolution using CIBERSORT (24). The signature matrix accurately classified the composition of flow-sorted OPCs, astrocytes, neurons, myelinating oligodendrocytes, endothelial cells, and microglia (Supplementary Fig. S3A and S3B). When applied to 150 dpi control samples, the inferred cell composition was overwhelmingly OPC with minor contributions of astrocytes and neurons, fragments of which likely wrapped around cells targeted by LCM (**Fig. 2C**). We considered *Cspg4-expressing* pericytes as an alternative source of contamination and substituted a set of pericyte markers (26) in place of the endothelial signature (Supplementary Fig. S3C and S3D). The inferred pericyte fraction was negligible, and the other cell proportions were similar to the first deconvolution but more variable as a result of pooling marker sets from different studies. We obtained similar results with 12 dpi and 90 dpi control samples (Supplementary Fig. S3E–H), supporting that cell purity does not change with tissue age. Together with the near-instantaneous cryopreservation (less than two minutes from sacrifice to embedding), the results bolster LCM as an effective means for labeled OPC isolation and transcriptomic profiling.

**Figure 2.**
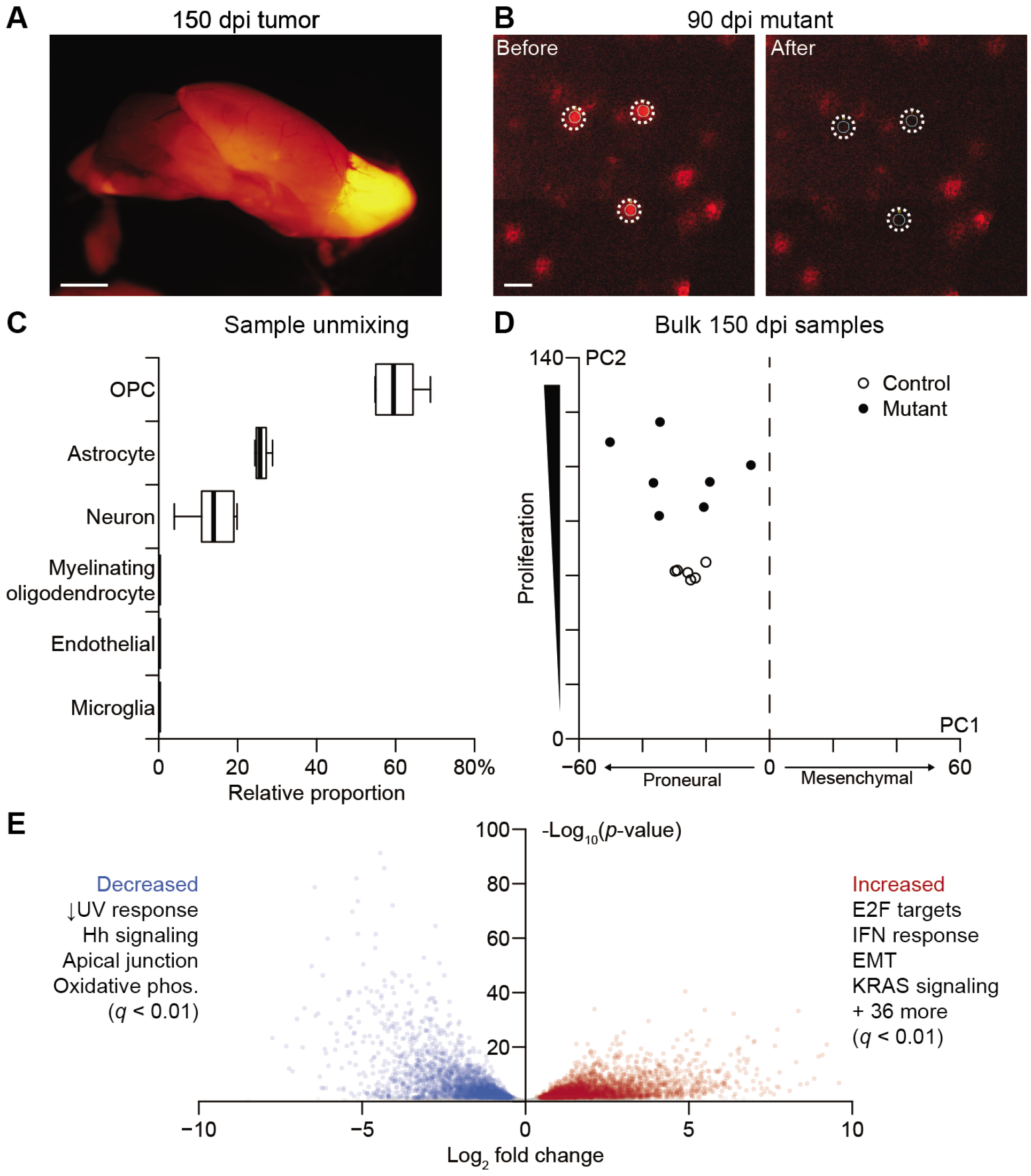
Bulk LCM transcriptomes of labeled gliomas are specific to OPC-derived cells and show gene-expression changes consistent with *Nf1–Trp53* deletion. **A,** Macroscopic image of a labeled glioma in the olfactory bulb of a 150 dpi animal. Scale bar is 2 mm. **B,** Microscopic image of 90 dpi mutant OPCs. Images were taken before and after LCM to show spatial fidelity of the isolation. Scale bar is 25 μm. **C,** Relative proportion of the indicated cell types in the control bulk RNA-seq samples estimated by CIBERSORT (24). Boxplots show the median and interquartile range with whiskers indicating the range from *n* = 6 independent samples of control 150 dpi OPCs. **D,** Projection of the bulk 150 dpi mutant and control samples onto principal components capturing proneural–mesenchymal state and relative proliferative status (3). **E,** Volcano plot of differentially expressed genes between tumors and controls. Enriched hallmark pathways among increased and decreased transcripts are summarized, and the complete list is available in Supplementary File S1.

We sequenced seven samples from tumors and six samples of OPCs from age-matched control mice, detecting 11,748 ± 651 genes at greater than five transcripts per million (TPM). To revisit the molecular subtype of glioma cells in bulk, we used a principal components-based projection that stratifies cells along two axes: proneural-vs.-mesenchymal differentiation and quiescent-vs.-proliferative activity (3). Both control OPCs and mutant tumors clearly resided on the proneural side of the projection, but there was greater variability observed among the tumors (**Fig. 2D**). As expected, tumors were also consistently more proliferative according to their projections along the second axis. We examined quantitative differences in transcript abundance between the two groups with DESeq2 (35), incorporating sex as a covariate (see Materials and Methods). The analysis identified 6057 differentially abundant transcripts (Supplementary File S2); there was a bias toward upregulation, consistent with the increased transcription driven by Ras signaling (3453 genes vs. 2604 genes, *p* < 10^-15^ by binomial test; **Fig. 2E**) (36). Neither *Trp53* nor *Nf1* total transcript counts were reduced based on RNA-seq reads, but deletion of the targeted exons was confirmed in the read distribution of the mutant alignments (Supplementary Fig. S4). In search of coherent pathways, we queried the Molecular Signatures Database (32), finding hallmarks related to *Trp53* loss (DNA damage), *Nf1* loss (Ras signaling, proliferation), and immune responses consistent with local activation of microglia (2) (**Fig. 2E** and Supplementary File S1). There were multiple increased transcripts for hallmarks of epithelial-to-mesenchymal transition (EMT)—*Bgn, Cd44, Mgp, Timp1, Tnc,* and *Vim* all increased with log_2_ fold change >3—but these changes were insufficient to displace the overall proneural projection of the tumors. The bulk-profiling results from 150 dpi samples validate the LCM procedure and reinforce that this GEMM gives rise to proneural gliomas with geneexpression patterns reflecting *Nf1–Trp53* loss.

### Transcript abundance changes detected by bulk analysis and cell-state heterogeneities detected by 10cRNA-seq are uncoupled in early mutant OPCs

We anticipated that the early responses to *Nf1–Trp53* loss would be more informative for tumorigenesis and more variable among labeled cells. Therefore, we coupled bulk transcriptomics with heterogeneity analysis by stochastic profiling (17,19) at 12 dpi to capture the immediate response of *Nf1–Trp53* deletion (**Fig. 1**). The two-pronged approach would enable a comparative assessment and provide complementary perspectives on the single-cell changes elicited by a two-hit tumor-suppressor loss in OPCs.

For the seven mutant OPC samples and five controls measured in bulk, we found that overall transcript content was very similar to that of the 150 dpi samples (11,674 ± 150 genes detected at greater than five TPM). Yet, when the 12 dpi bulk samples were projected onto the earlier principal-component axes (3), mutant and control OPCs overlapped as a tight cluster, all with moderate proliferation and a proneural phenotype (**Fig. 3A**). The result was surprising, because we previously reported that mutant OPCs incorporate BrdU at an elevated rate during the week after tamoxifen addition (15). The lack of a distinguishing RNA-seq projection here could be reconciled if the proliferative burst of the GEMM occurred immediately after *Nf1–Trp53* deletion was induced and then subsided by 12 dpi. Corroborating this notion, only 3.7% of tdTomato^+^ cells were positive for phosphorylated Rb or Ki67 in 12 dpi tissue sections, compared to 89% in 150 dpi samples (Supplementary Fig. S5).

**Figure 3.**
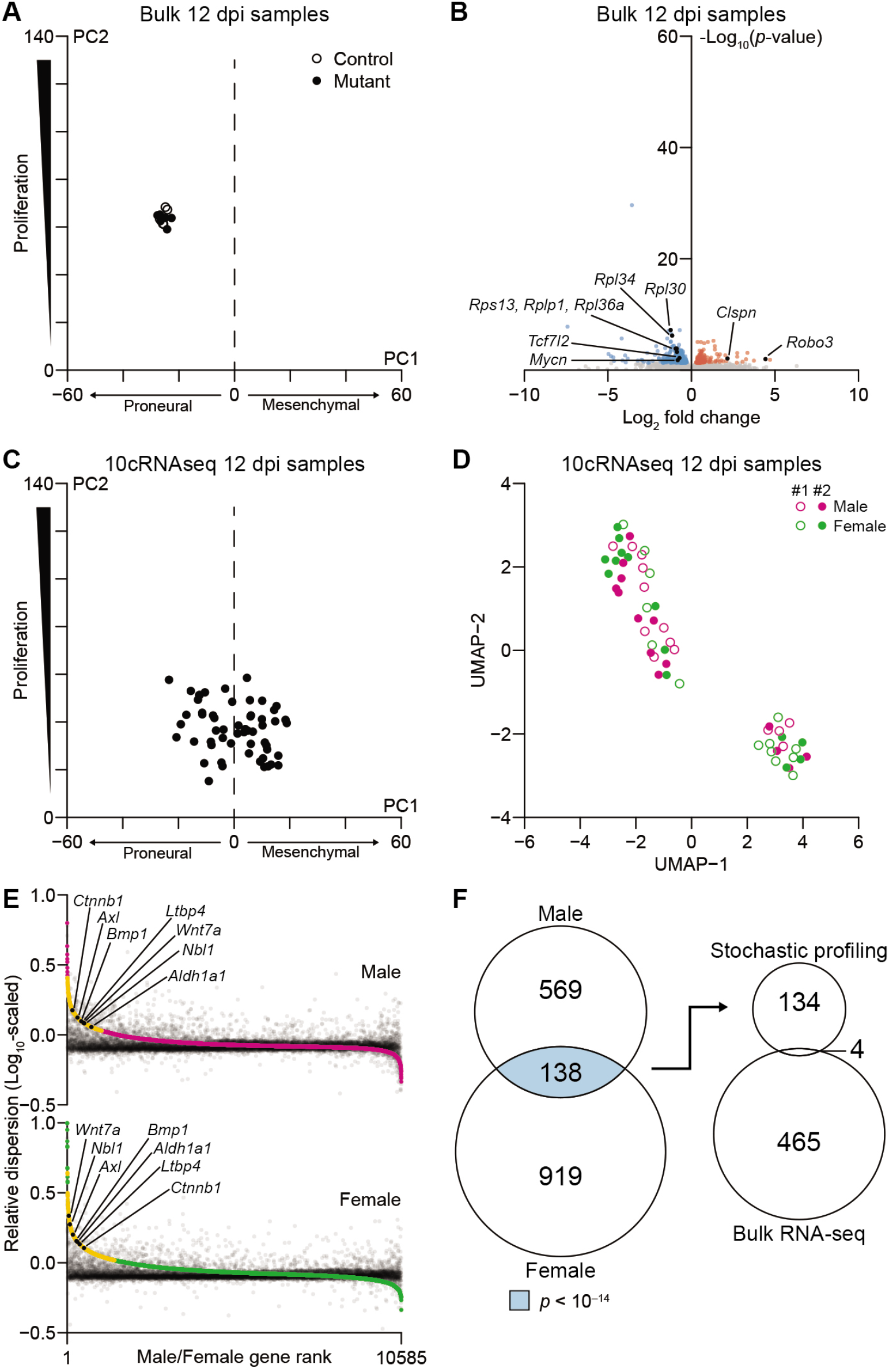
Bulk and 10-cell transcriptomics give orthogonal perspectives on the early adaptations of mutant cells to *Nf1–Trp53* deletion. **A,** Projection of the bulk 12 dpi mutant and control samples onto principal components capturing proneural–mesenchymal state and relative proliferative status (3). **B,** Volcano plot of differentially expressed genes between 12 dpi mutant samples and controls. Enriched hallmark pathways among increased and decreased transcripts are shown along with specific genes discussed in the text. Complete lists are available in Supplementary Files S1 and S2. **C,** Projection of the 10cRNA-seq 12 dpi mutant samples onto principal components capturing proneural–mesenchymal state and relative proliferative status (3). **D,** UMAP projection of 10cRNA-seq observations labeled by sex and mouse sample number. **E,** Abundance-dependence dispersion of pool-and-split controls (black) and 10-cell samples from males (magenta) and females (green). Significantly overdispersed genes for each sex are indicated in yellow, and genes discussed in the text are indicated in black. **F,** Venn diagram (left) intersecting male and female candidates to yield RHEGs (light blue). The overlap with differentially abundant genes (right) was not statistically significant (*p* = 0.53).

We performed a differential-expression analysis on the bulk samples and identified 469 transcripts elevated or reduced in mutant OPCs compared to control, 204 of which were identified at 150 dpi (*p* < 10^-15^ by hypergeometric test; **Fig. 2E, 3B**, and Supplementary File S2). Mutant OPCs showed large increases in the axon guidance receptor *Robo3* (37) (log_2_ fold change = 4.4) and *Clspn* (log_2_ fold change = 2.2), a scaffold coordinating the DNA replication checkpoint (38) (**Fig. 3B** and Supplementary File S2). Many ribosomal subunits were reciprocally decreased, likely due to the concomitant reduction of *Mycn* (39) (log_2_ fold change = –0.82) downstream of reduced *Tcf7l2* (40) (log_2_ fold change = –0.72). Unexpectedly, there was not an obvious coordination in expression changes stemming from recognized pathways. Hallmark gene-set enrichments were absent from the 12 dpi data except for the mitotic spindle, which among all increased transcripts was enriched just beyond the false-discovery rate (*q* < 0.01; Supplementary File S1). Our observations at 12 dpi suggested that the bulk RNA-seq data were capturing generic adaptations to *Nf1* loss unencumbered by Trp53 checkpoints.

Although bulk transcriptomes were very consistent at 12 dpi (**Fig. 3A**), we reasoned that the state of individual OPCs during and shortly after *Nf1–Trp53* deletion would be different. To evaluate regulatory heterogeneities in mutant OPCs, we carefully devised an experimental plan to apply stochastic profiling by 10cRNA-seq at 12 dpi. First, random 10-cell sampling was restricted to labeled OPCs in the olfactory bulb, where gliomas are most likely to arise in the GEMM (15). Samples were excluded if there was evidence of pericyte contamination by qPCR (Supplementary Table ST1). Second, we hedged against mouse-to-mouse variations by allocating 28 random samples and 20 pool-and-split controls equally between olfactory-bulb hemispheres of two separate animals. The samples and controls were merged for analysis (19) and inspected afterwards for mouse-dependent batch effects. Statistically, we built upon the resampling approach of the accompanying contribution (20) by evaluating overdispersion stability through leave-one-out cross-validation (see Materials and Methods). Consistent regulatory heterogeneities were readily delineated as those appearing in >75% of crossvalidation runs (Supplementary Fig. S6). Finally, given the sex differences documented in human GBM (33), the entire stochastic-profiling study was performed once in males and once in females. Intersecting the male and female candidates at the 12 dpi time point led to a set of recurrent, heterogeneously expressed genes (RHEGs) for gleaning cell states.

Before fluctuation analysis, we projected the 10cRNA-seq observations onto the principal-component axes (3) used previously with bulk data (**Fig. 2D** and **3A**). Proliferation scores were reduced and more variable (**Fig. 3C**), as expected from the ~100-fold reduction in cellular material. Projections on the proneural-to-mesenchymal spectrum were also displaced because 10-cell observations did not detect the strongest proneural markers as reliably as bulk RNA-seq and mesenchymal transcripts were sporadically more prominent (Supplementary Fig. S7). Furthermore, we found that the 10cRNA-seq samples were distinctly bimodal, half with more-proneural character and half with more-mesenchymal character (Supplementary Fig. S8A). These mixed mesenchymal profiles are consistent with those noted in the accompanying contribution among breast carcinoma cells (19). By contrast, predicted 10-cell distributions were exclusively proneural for normal OPCs drawn from two independent scRNA-seq datasets, indicating that mesenchymal projections were not intrinsic to the cell type [GSE60361 (29) and GSE75330 (30)] (Supplementary Fig. S8B and S8C). The observations in mutant OPCs were consistent with inferred single-cell profiles (23) that were more frequently mesenchymal than proneural compared to normal OPCs (see Materials and Methods; Supplementary Fig. S8D–F). However, the mesenchymal bias accentuated by 10-cell data reverted to the mean and disappeared when samples were agglomerated computationally as bulk averages of 700–1000 cells (Supplementary Fig. S8G). We independently probed the characteristics of mutant cells at 12 dpi by immunostaining for mesenchymal markers loaded strongly on the principal-component axis (Supplementary Fig. S9). Three of five targets showed positive immunoreactivity in a minor-but-significant fraction of mutant cells: Tgfbi (26%), Fn1 (23%), and Mcl1 (7.7%; all *p* << 10^-16^ by binomial test). Such expression frequencies are readily detectable as fluctuations in 10-cell samples (17,23), but their contribution to bulk profiles would be much more dilute. This first-pass comparison of measurement types indicated that 10cRNA-seq data had revealed early mesenchymal heterogeneity at the 12 dpi time point that was obscured in bulk observations.

Overall, across 56 10-cell samples of mutant OPCs (28 male + 28 female), we detected 4570 ± 1900 genes at greater than five TPM, suggesting that ~40% of the bulk transcriptome would be amenable for stochastic-profiling analysis. By comparison, applying a similar threshold to scRNA-seq data from flow-sorted OPCs (30) yielded ~15% fewer transcripts (3852 ± 956 genes, *p* < 0.01 by K-S test). We inspected the 10-cell data for confounders by using uniform manifold approximation and projection (UMAP) (41) to reduce dimensionality of the 10cRNA-seq profiles. Along two UMAP dimensions, there were 2–3 clusters of gene expression, each populated with 10-cell observations from different tissue batches and sexes at 12 dpi (**Fig. 3D**). The global projections illustrated that confounding batch effects were negligible and that cell-state heterogeneities might be embedded in the 10-cell transcript fluctuations from sample to sample.

Regulatory heterogeneities were assessed formally through abundance-dependent dispersion modeling (42,43) that used pool-and-split controls of 10-cell equivalents to exclude technical artifacts [see the accompanying contribution (19) for details]. High-ranking candidates showed 10-cell sampling fluctuations much greater than expected given their abundance and technical noise (**Fig. 3E**). At 12 dpi, we identified 707 candidate regulatory heterogeneities in males and 1057 in females (Supplementary File S3). The significantly higher proportion of candidates in females (*p* << 10^-16^ by sign-rank test) coincided with a modest enrichment for human GBM transcripts specific to females (*p* < 0.05 by hypergeometric test), who have a reduced GBM incidence and longer survival compared to males (33). We observed no such enrichment for male-specific transcripts among male regulatory heterogeneities, supporting a recent report that OPC diversity may be less pronounced in males (44). Importantly, between sexes, there was a highly significant enrichment of 138 RHEGs shared at 12 dpi (*p* < 10^-14^ by hypergeometric test; **Fig. 3F**, left, and Supplementary Fig. S10). Yet, only four of the RHEGs identified by stochastic profiling overlapped with the transcripts found to be differentially expressed in bulk 12 dpi samples (*p* = 0.53 by hypergeometric test; **Fig. 3F**, right). The intersections increased when data were stratified by sex first, but the number of overlapping genes remained insignificant (Supplementary Fig. S11A). We interpreted 12 dpi RHEGs as sex-independent variations in mutant cells that are almost entirely obscured at the population level.

There were multiple transcripts of interest in the 12 dpi RHEG set (Supplementary File S3). The receptor tyrosine kinase *Axl,* for instance, is implicated in oligodendrocyte survival and myelination after injury (45) and promotes self-renewal of mesenchymal progenitors in GBM (46). The stem-cell marker *Aldh1a1* was also a RHEG with links to an OPC subpopulation found at roughly equal proportions in males and females (44). Relevant to the mixed proneural– mesenchymal states (**Fig. 3C**), we found extensive evidence for heterogeneous regulation of transforming growth factor β (TGFβ)-family signaling (47,48). The latent TGFβ-binding protein *Ltbp4,* the TGFβ-superfamily ligand *Bmp1,* and the Bmp-subfamily inhibitor *Nbl1* (49) were all RHEGs (Supplementary File S3). Using *Nbl1* as a reference, we searched the 10cRNA-seq transcriptomes for strong covariates that fell below the statistical thresholds of the dispersion test (27), identifying the TGFβ-induced transcripts, *Snai2 (R* = 0.58) and *Vip (R* = 0.52) (50). Regulators of OPC state transitions were also prevalent RHEGs, including *Wnt7a* and *Ctnnb1* (51). Although some 12 dpi RHEGs reflect normal asynchronies in OPC maturation, they nonetheless give clues about different cellular contexts shortly after *Nf1–Trp53* loss.

### Premalignant glioma is punctuated by aberrant differentiation and a neomorphic state coupling homology-dependent repair and nonsense-mediated decay with DNA damage

The 12 dpi studies documented the initial OPC cell states and early adaptations to *Nf1– Trp53* loss. We suspected a very different set of transcriptional changes and variation at an intermediate time, when unproductive mutants have stalled or been culled and the remaining expansion of mutant OPCs adapts to premalignancy (**Fig. 1A**). For transcriptomic profiling, we selected 90 dpi, a time point when mutant OPC proliferation has yet to restart in the GEMM and the overall number of labeled cells remains stable (15). Even though tdTomato^+^ cells were more numerous in the olfactory bulb at 90 dpi, there is no evidence for local expansion of mutant clones (**Fig. 1A**) (15). Therefore, the study design for bulk and 10-cell sample acquisition was identical to 12 dpi to enable comparisons between time points.

The bulk 90 dpi profiling of seven mutant OPC samples and four controls yielded 11,577 ± 106 genes detected at greater than five TPM, very similar to the other time points. Despite months of evolution, the projection of bulk 90 dpi samples onto proliferative and proneural– mesenchymal principal components was indistinguishable from their 12 dpi counterparts (**Fig. 3A** and **4A**). The subsequent divergence of 150 dpi gliomas on these principal components (**Fig. 2D**) suggests an even-later event is needed to drive malignant transformation in the GEMM. Despite the increased time for adaptation at 90 dpi, there were significantly fewer differentially expressed genes than at 12 dpi, with only 82 increased and 61 decreased (*p* < 10^-15^ by binomial test; **Fig. 4B**). Although there was no hallmark gene-set enrichment among the upregulated transcripts, we noted a few DNA-replication markers—*Top2a, Ccna2, Hjurp—*that increased slightly with log_2_ fold change = 1.2–1.9 (**Fig. 4B** and Supplementary File S2). Thus, despite undetectable increases in BrdU incorporation at 90 dpi (15) and lack of an overarching proliferative signature (**Fig. 4A**), more mutant OPCs appeared to be entering S phase than control. Reduced transcripts included many high-abundance markers for pericytes (*Pdgfrb, Cd248, Anpep*) (26), suggesting that their minor contamination had been further diluted out by the expanded mutant OPCs.

**Figure 4.**
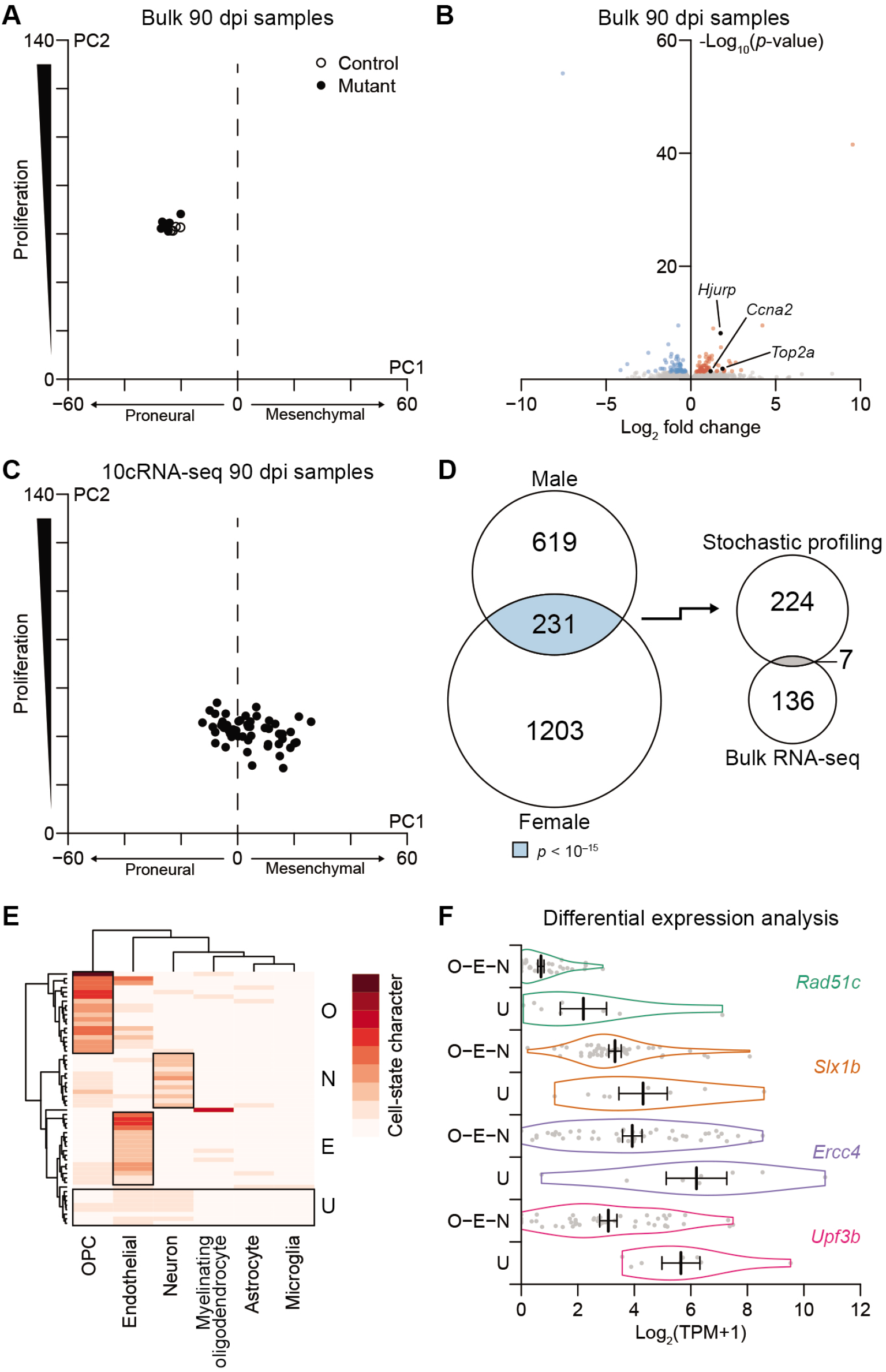
Premalignant mutants exhibit few population-level changes in transcript abundance despite gross alterations in single-cell fate and state profiles. **A,** Projection of the bulk 90 dpi mutant and control samples onto principal components capturing proneural–mesenchymal state and relative proliferative status (3). **B,** Volcano plot of differentially expressed genes between 90 dpi mutant samples and controls. Enriched hallmark pathways among increased and decreased transcripts are shown along with specific genes discussed in the text. Complete lists are available in Supplementary Files S1 and S2. **C,** Projection of the 10cRNA-seq 12 dpi mutant samples onto principal components capturing proneural–mesenchymal state and relative proliferative status (3). **D,** Venn diagram (left) intersecting male and female candidates to yield RHEGs (light blue). The overlap with differentially abundant genes (right) was marginally significant (gray, *p* < 0.05). **E,** Absolute cell-state character for the indicated marker sets in the 10cRNA-seq samples estimated by CIBERSORT (24). Sample groups with substantial cell-state character of OPCs (O), endothelial cells (E), or neurons (N) are marked along with an undefined group (U). **F,** Violin plots for a subset of transcripts significantly increased in the U group compared to the O–E–N groups. Data are shown as the mean ± s.e.m. of *n* = 46 (O–E–N) and 8 (U) 10-cell observations at 90 dpi.

The projections of 10cRNA-seq data collected at 90 dpi (5649 ± 1197 genes at greater than five TPM) were likewise comparable to those at 12 dpi, except that the range of proliferation scores was 1.5-fold smaller (**Fig. 3C** and **4C**), indicating a narrowed breadth of cells in a cycling-like state. Additionally, we found that cell-state scores became sexually dimorphic, with female samples residing on the proneural end of the spectrum and males on the mesenchymal end (Supplementary Fig. S12). Intersection of male and female candidate heterogeneities identified 231 RHEGs (*p* < 10^-15^ by hypergeometric test), a significant increase compared to the 138 RHEGs at 12 dpi (*p* < 10^-12^ by binomial test; **Fig. 3F** and **4D**, Supplementary Fig. S13, and Supplementary File S3) and a significant increase compared to the bulk analysis at 90 dpi (*p* < 10^-11^ by binomial test). As before, very few RHEGs were differentially abundant in bulk samples and vice versa (**Fig. 4D** and Supplementary Fig. S11B), reiterating the information gained by stochastic profiling over the bulk approach.

The larger number of RHEGs (*p* < 10^-10^ by binomial test) suggested that a deeper investigation of cell-state heterogeneity at 90 dpi would be informative. The RHEG *Ly6a* (a murine stem-cell marker) covaried with the reprogramming factor *Klf4 (ρ* = 0.54) and the mesenchymal transcript *Vim (ρ* = 0.41) that lay in the union set of candidates (**Fig. 4D**). Further, there were various markers for oligodendrocytes (*Mog, Cldn11, Opalin, Mbp*) as reported in human gliomas (9), neurons (*Syt7, Resp18, Stmn2*) as reported in human GBMs (5), and pericyte-like cells (*Rgs5, Vtn*) (52) within the 90 dpi RHEG set (Supplementary File S3). In bulk samples, the 90 dpi time point showed no differences in overall purity compared to 12 dpi or 150 dpi (Supplementary Fig. S3C–H). We interpreted the mixed distribution of fate markers as a chaotic (de)differentiation of mutant OPCs into states partly reflecting other cells of the brain.

To quantify cell-state character, we returned to CIBERSORT, applying the algorithm in absolute mode for estimating how much of each 10-cell profile was attributable to markers in the signature matrix (**Fig. 4E** and Supplemental Fig. S3A). 34% of 90 dpi observations retained OPC as the most dominant contributing signature (O group, **Fig. 4E**). Observations with strong endothelial contributions suggested a confused mural-cell state (52) (E group, **Fig. 4E**). We also found 18% of samples with dominant (albeit weaker) contributions from neuronal signatures (N group, **Fig. 4E**). Neuron-like cells are absent in various subtypes of glioma (7–9), suggesting the N group may contain cellular dead-ends for gliomagenesis. Consistent with prior immunocytochemistry (15), samples with properly differentiated myelinating oligodendrocytes (MO) were rare. However, when we applied absolute CIBERSORT to 10-cell profiles at 12 dpi, there was a considerable MO group (Supplementary Fig. S14A), suggesting that a third deadend subpopulation had already been eliminated by death or dedifferentiation at 90 dpi. Most importantly, we identified 14% of 10-cell samples lacking strong contributions from any conventional cell type in the brain, marking the group as undefined (U group, **Fig. 4E**). The U group was specific to 90 dpi, as there were no such profiles in 10-cell samples at 12 dpi or in bulk profles at 12, 90, or 150 dpi (Supplementary Fig. S14A,B). The results indicated that bulk tumor deconvolution of 10cRNA-seq pools could identify novel cell states warranting further analysis.

We hypothesized that the mysterious U group would contain information about a premalignant bottleneck. When the U group was separated from the O–E–N samples and evaluated for differential transcripts (see Materials and Methods), we found widespread downregulation of 163 genes. Among the 13 transcripts with increased abundance in the U group were *Upf3b,* an X-linked regulator of nonsense-mediated decay (53), and a set of genes directly involved in resolving Holliday junctions during homology-dependent repair: *Rad51c, Slx1b* (the mouse ortholog of human *SLX1),* and *Ercc4* (54,55) (**Fig. 4F**). The U group also showed an increase in *Bicra,* a glioma tumor-suppressor candidate with a human polymorphism that predisposes for 19q deletions in oligodendroglioma (56). None of these transcripts were differentially expressed in bulk 150 dpi tumor samples compared to control (log_2_ fold change: – 0.25 to 0.32, *q* > 0.4; Supplementary File S1), suggesting a transient cell state in a minority of the 90-dpi mutant population.

The Rad51 and Upf families interact genetically in yeast during homology-dependent repair after DNA damage (57). We hypothesized that a similar coordination might occur with mutant cells in the U state and immunostained 90-dpi cryosections for a marker of DNA damage (γH2AX—phospho-H2A.X Ser^139^) together with either Ercc4 or Upf3b. Among tdTomato^+^ mutant cells with high Ercc4 (~25%) or Upf3b (~16%), there was a significant increase in γH2AX immunoreactivity (**Fig. 5A–F**), suggesting DNA damage as a U-state trigger. We confirmed the 10cRNA-seq-predicted coupling of the U state by staining cells for Ercc4 and Upf3b concurrently, finding that the two proteins were mutually co-expressed (**Fig. 5G–I**). Without *Trp53* function, homology-dependent repair and nonsense-mediated decay appeared to create a regulatory choke point of persistent DNA damage, which could blur cell identity and lead to chromosomal instability during late premalignancy.

**Figure 5.**
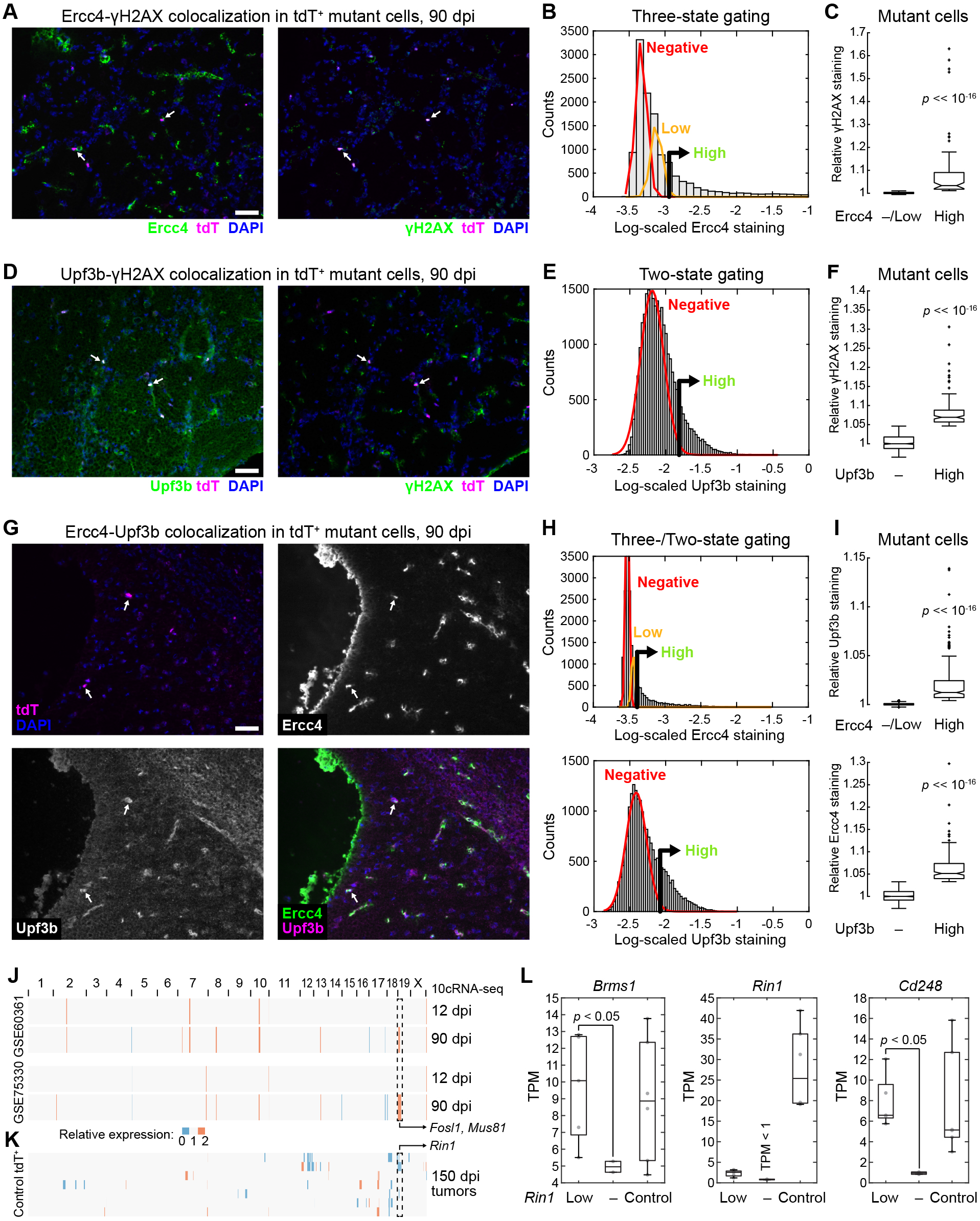
Mutant cells sporadically co-express regulators of homology-dependent repair and nonsense-mediated decay coincident with a marker of DNA damage. **A-F,** Representative images (**A** and **D**), quantitative immunocytochemistry (**B** and **E**), and grouped comparisons (**C** and **F**) of 90 dpi cryosections stained concurrently for Ercc4 (**A-C**) or Upf3b (**D-F**) and phospho-H2A.X Ser^139^ (γH2AX). Summary quantifications are from 12,366 (**B**) or 26,556 (**E**) nuclei (gray) and 176 (**C**) or 915 (**F**) tdTomato^+^ mutant cells from *n* = 4 tissues of three animals. **G-I,** Representative multichannel images (**G**), quantitative immunocytochemistry (**H**), and grouped comparisons (**I**) of 90 dpi cryosections stained concurrently for Ercc4 and Upf3b. Summary quantifications are from 19,865 nuclei and 1116 tdTomato^+^ mutant cells from *n* = 4 tissues of three animals. (**J** and **K**) inferCNV profiles estimating mean-averaged regional gene expression in 12 and 90 dpi 10cRNA-seq samples (**J**) or individual chromosomal gains–losses in 150 dpi bulk tumors (**K**). Reference transcriptomes used are shown to the left. (**L**) Loss of *Rin1* detection coincides with significant reductions in its flanking genes, *Brms1* and *Cd248.* Data are shown as boxplots of *n* = 2–6 animals from each group. Differences were assessed by one-way rank sum test. In **B, E**, and **H**, positive gates were set at the 99^th^ percentile of the low (Ercc4) or absent (Upf3b) subpopulation. High vs. Negative (–)/Low subpopulations were compared by rank sum test.

Considering the S-phase markers and widespread transcriptional variation (**Fig. 4B,D**), we asked whether 90-dpi mutant cells might be experiencing transcription-induced replication stress. Activated Ras (i.e., from *Nf1* deletion) elevates overall transcriptional activity, RNA polymerase occupancy, and the formation of RNA–DNA hybrids (“R loops”), which interfere with the replication machinery and can give rise to DNA damage (36). Our hypothesis was that loci of elevated transcription would have enhanced R loops and DNA damage that result in selective chromosomal alterations later in gliomagenesis. We used inferCNV to localize regionally elevated transcript expression to specific genomic coordinates. For two normal OPC reference transcriptomes, we identified a ~2.4 Mb segment on mouse chromosome 19 (mChr19) that became elevated specifically in 90-dpi 10cRNA-seq samples (**Fig. 5J** and Supplementary Fig. S15A). Within this locus reside *Fosl1* (58), a recently reported driver of the mesenchymal transition of *Nf1-null* glioblastoma, and *Mus81,* an endonuclease that is critical for resolving R loops (59). Mus81 also resides in the same Holliday-junction repair complex as Slx1b and Ercc4 (55), further supporting why the locus might selectively require hyperactivation at 90 dpi.

The elevated inferCNV signals at 90 dpi were not chromosomal gains, because there was no evidence of amplification in bulk transcriptomes from 150-dpi tumors by the same measure (**Fig. 5K** and Supplementary Fig. S15B). In fact, among 150-dpi tumors, we identified multiple instances of inferred loss in the same mChr19 locus, suggesting an important restraint to gliomagenesis was located in the actively transcribed region. After cross-referencing with bulk differential expression (Supplementary File S2), we identified the Ras interactor, *Rin1.* Biochemically, Rin1 binds tightly to Ras at the expense of Raf1 and downstream signaling through MEK–ERK (60). Thus, loss of Rin1 should promote ERK signaling from Ras activated by the loss of *Nf1*. Whereas *Rin1* is reduced in all 150-dpi tumors, supporting the need for its removal, only in cases where *Rin1* was “absent” (TPM < 1) did we find coincident downregulation in its nearby flanking genes, *Brms1* and *Cd248* (*p* < 0.05 by one-sided rank-sum test; **Fig. 5L**). Although *Rin1* abundance is clearly reduced by multiple mechanisms, sustained upregulation of transcription-coupled endonucleases *Ercc4* and *Mus81* coincides with microdeletion.

### Integrating bulk and 10-cell transcriptomes defines self-consistent trajectories of gliomagenesis

Differential transcripts and RHEGs showed little overlap at 12 dpi and 90 dpi (**Fig. 3F** and **4D**), raising the possibility that the data types were too dissimilar for meaningful comparison. We asked whether the bulk and 10cRNA-seq data could be reconciled by fusing the observations according to only those transcripts that were differentially abundant or recurrently heterogeneous at least once (see Materials and Methods). The datasets were fused after standardization and reduced to two UMAP dimensions, with samples displayed as subsets for inspection of the median centroids pertaining to each group. The goal was to evaluate bulk and 10-cell sample trajectories on the UMAP for themes common to both data types.

With the bulk samples obtained at 150 dpi, we noted a diagonal trajectory from control OPC samples to tumor cells along the two UMAP dimensions (**Fig. 6A**). We interpreted this trajectory as a gliomagenesis “axis” within the UMAP. For 12 dpi samples, the trajectory from control to mutant was nearly perpendicular to that at 150 dpi (**Fig. 6B**) and delineated an adaptive axis, which was distinct from the late readouts of tumor-suppressor loss projected by the 150 dpi tumors. Most interesting were the 90 dpi trajectories, in which bulk mutant samples appeared to bifurcate in two directions from the controls (**Fig. 6C**). Half of the 90 dpi mutant samples were similar to 12 dpi bulk observations, indicating an averaged population that was still undergoing adaptation, while the other half mapped toward the 150 dpi tumors, suggesting a fraction of cells in those bulk samples were transitioning to a gliomagenic state. The confused (de)differentiation of mutant OPCs during premalignancy [conceptually similar to that in the accompanying contribution (20)] undoubtedly contributed intrinsic variance to the very small number of differential transcripts detected at 90 dpi (**Fig. 4B**).

**Figure 6.**
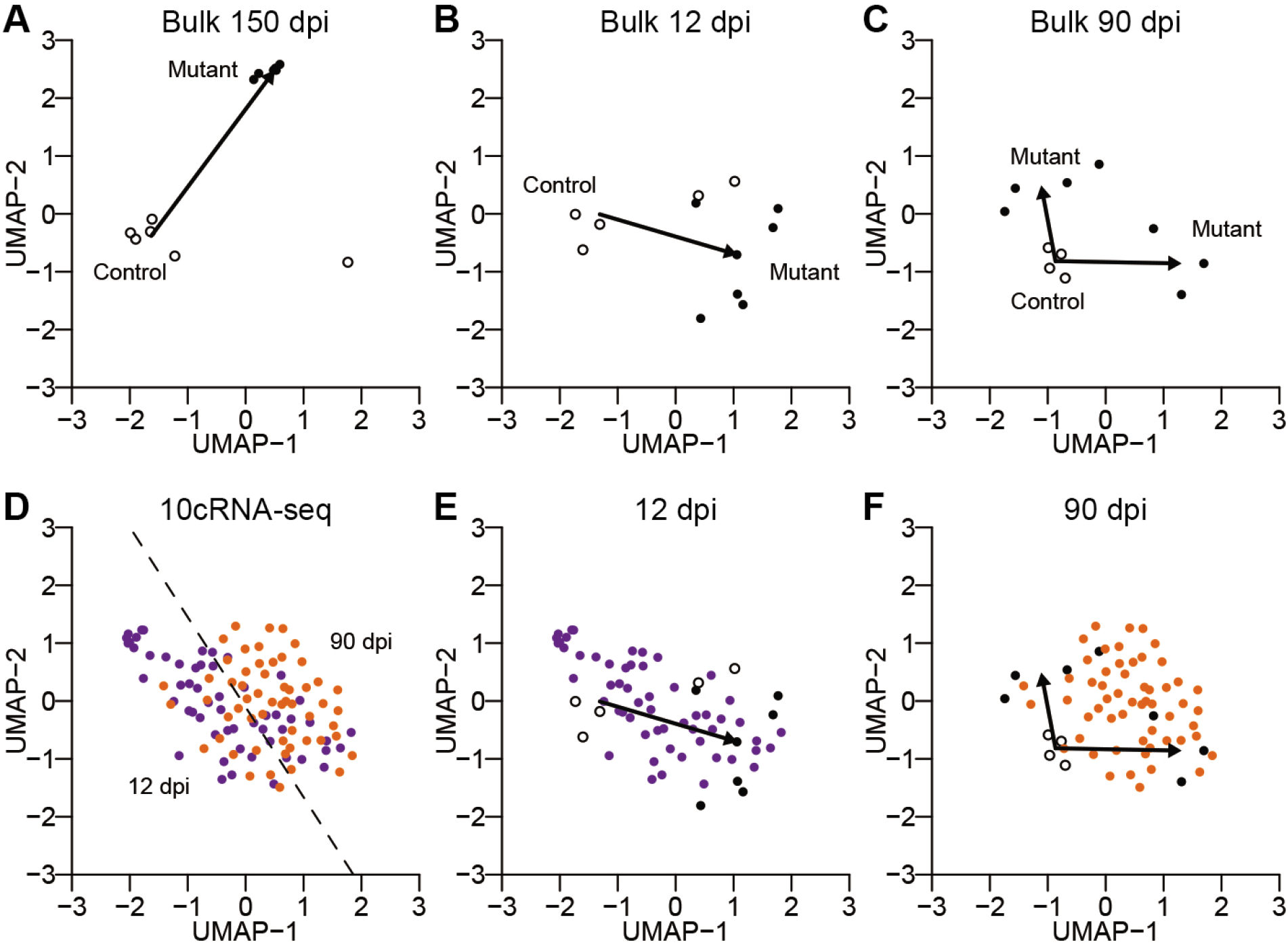
An integrated map of mutant gliomagenesis at the bulk and subpopulation levels. **A– C,** UMAP visualization of control and mutant bulk samples at 150 dpi (**A**), 12 dpi (**B**), and 90 dpi (**C**). Arrows indicate the vector of the control median centroid to the mutant median centroid(s). Two mutant centroids (identified by k-means clustering with two groups) were used at 90 dpi. **D–F,** UMAP visualization of 10cRNA-seq data at 12 dpi (purple) and 90 dpi (orange). The 12 dpi–90 dpi-separating hyperplane (identified by a support vector machine) is shown in **D**. Bulk and 10cRNA-seq samples are replotted together at 12 dpi (**E**) and 90 dpi (**F**). A single UMAP was generated using all of the data in **Fig. 6**, with indicated data subsets shown in individual subpanels.

Finally, we visualized the 10cRNA-seq data with the same UMAP cartography (**Fig. 6D**). The 12 dpi samples formed a cloud of observations that resided nearby controls and bulk mutant OPC observations at 12 dpi (**Fig. 6D** and **6E**), consistent with variable extents of progressive adaptation among mutant OPCs. In contrast, the 10-cell samples at 90 dpi were spanned by the bifurcating 90 dpi mutant OPC profiles measured in bulk (**Fig. 6D** and **6F**). We conclude that the 10cRNA-seq observations here reflect piecewise instances of premalignancy that contribute to the extreme variability in mutant OPC cell states before glioma onset. Thus, bulk and 10-cell data can be reconciled when viewed through their aggregate transcripts of interest.

## Discussion

Modern studies of intratumor heterogeneity strive for robust assessments of differential expression at single-cell resolution (5,7,9). For bulk tumors, quantitative changes in transcript abundance are often explained by differences in cell-type composition (8). Our study goes to the next level of granularity by examining bulk vs. single-cell differences in one cell type marked with a genetically defined perturbation that predisposes for glioma. The heterogeneity analysis uncovered differences in proneural–mesenchymal states and cell-fate assignments that were obscured in bulk averages. However, the 10-cell profiles also averaged over most cycling transcripts (19), many of which were detectably increased in bulk mutant OPCs compared to control. Other transcripts with bulk changes in expression did not fluctuate any more than expected given their measured abundance, implying that these alterations must occur somewhat uniformly in mutant OPCs. Amidst a relatively homogeneous adaptation to *Nf1– Trp53* deletion, stochastic events relating to OPC differentiation and replication stress recur until a productive mutant emerges (15).

For both time points analyzed by stochastic profiling, we identified substantially more regulatory heterogeneities in mutant OPCs from females compared to males. The difference is not easily explained by asynchronous hormonal cycles, because the observed variability was shared across samples. We also did not find an enrichment for female-specific candidates on the X chromosome, excluding random X inactivation as a possible explanation. The many female-specific heterogeneities at 12 dpi suggest a larger starting diversity of OPCs. Indeed, a recent study indicates that two OPC subpopulations predominate in males, whereas three subpopulations exist at roughly equal proportion in females (44). In addition to less regulatory diversity, the male 10-cell projections at 90 dpi scored as much more mesenchymal than females. This observation could relate to disease progression, as *Nf1–Trp53-predisposed* males are more likely to acquire further tumor-suppressor losses related to the mesenchymal subtype (61). Conceivably, reduced cellular diversity in males could increase the chance of gliomagenic events occurring in a mesenchymally susceptible subpopulation of OPCs.

The 90 dpi samples were unmixed into an assortment of cell states, but what ultimately enables a premalignant mutant OPC to become one of the singular gliomas of the GEMM (15)? We surmise that mechanisms for gliomagenesis are embedded in the RHEG set at 90 dpi. Transcriptional variability may act as a preamble to genomic alterations, such as the driver mutations described in the accompanying contribution (19) and the *Rin1* deletion identified here. For example, oligodendroglial tumors are tied to a loss of asymmetric cell divisions by OPCs and their differentiated progeny (62). The asymmetry regulator *Trim3* (63) is a 90 dpi RHEG, and Notch ligands appear separately as candidates in males (*Notch1*) and females (*Notch3).* Notch signaling was an artifactual signal in the accompanying contribution on small-cell lung cancer (20), but it could be a bona fide regulatory-state heterogeneity in mutant OPCs. Another probable contributor is cell-cycle checkpoint resolution, as 90 dpi RHEGs contain many individual checkpoint regulators: *Bap1, Brcc3–Anapc7* (coexpressed), *Trp53bp1, Nek9.* Last, we leave open the possibility of entirely new regulatory states, which are specifically critical for glioma premalignancy. Around the stem-progenitor RHEG *Lgr6,* for instance, there is an intriguing group of co-clustering RHEGs that includes a survival kinase (*Pim2),* a neural-glial adhesion ligand (*Astn2),* a tumor-promoting lysine methyltransferase (*Smyd2),* and a protooncogene (*Mllt11*) (Supplementary Fig. S10). In the brain, murine *Lgr6* expression is restricted to the olfactory bulb (64), providing a hypothesis for why it is a “hotspot” of gliomagenesis in this GEMM.

Gliomas establish simpler, hierarchical ecosystems (7–9) compared to their more advanced GBM counterparts (5). However, our results with one GEMM here suggest that paths to glioma ecosystems are chaotic and wasteful. Despite premalignant expansion, mutant OPC clones remain sparsely distributed up until onset of just a few focal gliomas (15), implying that multiple evolutionary dead-ends must be avoided, perhaps simultaneously, to progress. The bulk and 10-cell transcriptomic resources here give a roadmap for identifying the critical bottlenecks of proneural gliomagenesis. Future work will investigate the persistence of the regulatory-state heterogeneities identified and the fate of particular mutant cells. It will also be of great interest to examine the generality of these themes in other GEMMs and in human gliomas (7–9). Understanding such regulatory choke points could give insight into strategies for early detection and cancer prevention.

## Acknowledgements

We thank Emily Farber and Suna Onengut-Gumuscu at the UVA Center for Public Health Genomics for RNA-seq library preparation and sequencing, Craig Rumpel at the UVA Biorepository and Tissue Research Facility for LCM maintenance and histology services, UVA Research Computing for high-performance computing access and consulting, Henry Pritchard for assistance with NCBI GEO deposition, and Benjamin Purow, Roger Abounader, and Cheryl Borgman for critical evaluation of the manuscript. This work was supported by the National Institutes of Health #R01-CA194470 (K.A.J.), #U01-CA215794 (K.A.J.), and #R01-CA136495 (H.Z.), the David & Lucile Packard Foundation #2009-34710 (K.A.J.), the W. M. Keck Foundation (H.Z.), a UVA Cancer Center support grant #P30-CA044579, the UVA Medical Scientist Training Program (S.S.), and a Wagner Fellowship (S.S.).

## Notes

The authors declare no potential conflicts of interest.

### Competing Interest Statement

The authors have declared no competing interest.

### Summary of Updates

Changed title and updated figures and supplements to address reviewer comments.

## References

1. Verhaak RG, Hoadley KA, Purdom E, Wang V, Qi Y, Wilkerson MD, et al. Integrated genomic analysis identifies clinically relevant subtypes of glioblastoma characterized by abnormalities in PDGFRA, IDH1, EGFR, and NF1. Cancer Cell 2010;17:98–110

2. Wang Q, Hu B, Hu X, Kim H, Squatrito M, Scarpace L, et al. Tumor Evolution of Glioma-Intrinsic Gene Expression Subtypes Associates with Immunological Changes in the Microenvironment. Cancer Cell 2017;32:42–56 e6

3. Wang L, Babikir H, Muller S, Yagnik G, Shamardani K, Catalan F, et al. The Phenotypes of Proliferating Glioblastoma Cells Reside on a Single Axis of Variation. Cancer Discov 2019;9:1708–19

4. Patel AP, Tirosh I, Trombetta JJ, Shalek AK, Gillespie SM, Wakimoto H, et al. Single-cell RNA-seq highlights intratumoral heterogeneity in primary glioblastoma. Science 2014;344:1396–401

5. Neftel C, Laffy J, Filbin MG, Hara T, Shore ME, Rahme GJ, et al. An Integrative Model of Cellular States, Plasticity, and Genetics for Glioblastoma. Cell 2019;178:835–49 e21

6. Yuan J, Levitin HM, Frattini V, Bush EC, Boyett DM, Samanamud J, et al. Single-cell transcriptome analysis of lineage diversity in high-grade glioma. Genome Med 2018;10:57

7. Tirosh I, Venteicher AS, Hebert C, Escalante LE, Patel AP, Yizhak K, et al. Single-cell RNA-seq supports a developmental hierarchy in human oligodendroglioma. Nature 2016;539:309–13

8. Venteicher AS, Tirosh I, Hebert C, Yizhak K, Neftel C, Filbin MG, et al. Decoupling genetics, lineages, and microenvironment in IDH-mutant gliomas by single-cell RNA-seq. Science 2017;355

9. Filbin MG, Tirosh I, Hovestadt V, Shaw ML, Escalante LE, Mathewson ND, et al. Developmental and oncogenic programs in H3K27M gliomas dissected by single-cell RNA-seq. Science 2018;360:331–5

10. Parsa CF, Hoyt CS, Lesser RL, Weinstein JM, Strother CM, Muci-Mendoza R, et al. Spontaneous regression of optic gliomas: thirteen cases documented by serial neuroimaging. Arch Ophthalmol 2001;119:516–29

11. Network TCGA. Comprehensive genomic characterization defines human glioblastoma genes and core pathways. Nature 2008;455:1061–8

12. Liu C, Sage JC, Miller MR, Verhaak RG, Hippenmeyer S, Vogel H, et al. Mosaic analysis with double markers reveals tumor cell of origin in glioma. Cell 2011;146:209–21

13. Persson AI, Petritsch C, Swartling FJ, Itsara M, Sim FJ, Auvergne R, et al. Non-stem cell origin for oligodendroglioma. Cancer Cell 2010;18:669–82

14. Ledur PF, Liu C, He H, Harris AR, Minussi DC, Zhou HY, et al. Culture conditions tailored to the cell of origin are critical for maintaining native properties and tumorigenicity of glioma cells. Neuro Oncol 2016;18:1413–24

15. Galvao RP, Kasina A, McNeill RS, Harbin JE, Foreman O, Verhaak RG, et al. Transformation of quiescent adult oligodendrocyte precursor cells into malignant glioma through a multistep reactivation process. Proc Natl Acad Sci U S A 2014;111:E4214–23

16. Gonzalez PP, Kim J, Galvao RP, Cruickshanks N, Abounader R, Zong H. p53 and NF 1 loss plays distinct but complementary roles in glioma initiation and progression. Glia 2018;66:999–1015

17. Janes KA, Wang CC, Holmberg KJ, Cabral K, Brugge JS. Identifying single-cell molecular programs by stochastic profiling. Nat Methods 2010;7:311–7

18. Singh S, Wang L, Schaff DL, Sutcliffe MD, Koeppel AF, Kim J, et al. In situ 10-cell RNA sequencing in tissue and tumor biopsy samples. Sci Rep 2019;9:4836

19. Singh S, Sutcliffe MD, Repich K, Atkins KA, Harvey J, Janes KA. Pan-cancer drivers are recurrent transcriptional regulatory heterogeneities in early-stage luminal breast cancer. Cancer Res 2020:co-submitted

20. Schaff DL, Singh S, Kim KB, Sutcliffe MD, Park KS, Janes KA. Fragmentation of smallcell lung cancer regulatory states in heterotypic microenvironments. Cancer Res 2020:co-submitted

21. Malhotra A, Lindberg M, Faust GG, Leibowitz ML, Clark RA, Layer RM, et al. Breakpoint profiling of 64 cancer genomes reveals numerous complex rearrangements spawned by homology-independent mechanisms. Genome Res 2013;23:762–76

22. Miller-Jensen K, Janes KA, Brugge JS, Lauffenburger DA. Common effector processing mediates cell-specific responses to stimuli. Nature 2007;448:604–8

23. Bajikar SS, Fuchs C, Roller A, Theis FJ, Janes KA. Parameterizing cell-to-cell regulatory heterogeneities via stochastic transcriptional profiles. Proc Natl Acad Sci U S A 2014;111:E626–35

24. Newman AM, Liu CL, Green MR, Gentles AJ, Feng W, Xu Y, et al. Robust enumeration of cell subsets from tissue expression profiles. Nat Methods 2015;12:453–7

25. Zhang Y, Chen K, Sloan SA, Bennett ML, Scholze AR, O’Keeffe S, et al. An RNA-sequencing transcriptome and splicing database of glia, neurons, and vascular cells of the cerebral cortex. J Neurosci 2014;34:11929–47

26. He L, Vanlandewijck M, Raschperger E, Andaloussi Mae M, Jung B, Lebouvier T, et al. Analysis of the brain mural cell transcriptome. Sci Rep 2016;6:35108

27. Pereira EJ, Burns JS, Lee CY, Marohl T, Calderon D, Wang L, et al. Sporadic activation of an oxidative stress-dependent NRF2-p53 signaling network in breast epithelial spheroids and premalignancies. Sci Signal 2020;13:eaba4200

28. McQuin C, Goodman A, Chernyshev V, Kamentsky L, Cimini BA, Karhohs KW, et al. CellProfiler 3.0: Next-generation image processing for biology. PLoS Biol 2018;16:e2005970

29. Zeisel A, Munoz-Manchado AB, Codeluppi S, Lonnerberg P, La Manno G, Jureus A, et al. Brain structure. Cell types in the mouse cortex and hippocampus revealed by single-cell RNA-seq. Science 2015;347:1138–42

30. Marques S, Zeisel A, Codeluppi S, van Bruggen D, Mendanha Falcao A, Xiao L, et al. Oligodendrocyte heterogeneity in the mouse juvenile and adult central nervous system. Science 2016;352:1326–9

31. Wang L, Janes KA. Stochastic profiling of transcriptional regulatory heterogeneities in tissues, tumors and cultured cells. Nat Protoc 2013;8:282–301

32. Liberzon A, Birger C, Thorvaldsdottir H, Ghandi M, Mesirov JP, Tamayo P. The Molecular Signatures Database (MSigDB) hallmark gene set collection. Cell Syst 2015;1:417–25

33. Yang W, Warrington NM, Taylor SJ, Whitmire P, Carrasco E, Singleton KW, et al. Sex differences in GBM revealed by analysis of patient imaging, transcriptome, and survival data. Sci Transl Med 2019;11

34. Yao M, Ventura PB, Jiang Y, Rodriguez FJ, Wang L, Perry JSA, et al. Astrocytic transDifferentiation Completes a Multicellular Paracrine Feedback Loop Required for Medulloblastoma Tumor Growth. Cell 2020;180:502–20 e19

35. Love MI, Huber W, Anders S. Moderated estimation of fold change and dispersion for RNA-seq data with DESeq2. Genome Biol 2014;15:550

36. Kotsantis P, Silva LM, Irmscher S, Jones RM, Folkes L, Gromak N, et al. Increased global transcription activity as a mechanism of replication stress in cancer. Nat Commun 2016;7:13087

37. Zelina P, Blockus H, Zagar Y, Peres A, Friocourt F, Wu Z, et al. Signaling switch of the axon guidance receptor Robo3 during vertebrate evolution. Neuron 2014;84:1258–72

38. Smits VAJ, Cabrera E, Freire R, Gillespie DA. Claspin - checkpoint adaptor and DNA replication factor. FEBS J 2019;286:441–55

39. Boon K, Caron HN, van Asperen R, Valentijn L, Hermus MC, van Sluis P, et al. N-myc enhances the expression of a large set of genes functioning in ribosome biogenesis and protein synthesis. EMBO J 2001;20:1383–93

40. Shu W, Guttentag S, Wang Z, Andl T, Ballard P, Lu MM, et al. Wnt/beta-catenin signaling acts upstream of N-myc, BMP4, and FGF signaling to regulate proximal-distal patterning in the lung. Dev Biol 2005;283:226–39

41. Becht E, McInnes L, Healy J, Dutertre CA, Kwok IWH, Ng LG, et al. Dimensionality reduction for visualizing single-cell data using UMAP. Nat Biotechnol 2018

42. Kharchenko PV, Silberstein L, Scadden DT. Bayesian approach to single-cell differential expression analysis. Nat Methods 2014;11:740–2

43. Fan J, Salathia N, Liu R, Kaeser GE, Yung YC, Herman JL, et al. Characterizing transcriptional heterogeneity through pathway and gene set overdispersion analysis. Nat Methods 2016;13:241–4

44. Beiter RM, Fernández-Castañeda A, Rivet-Noor C, Merchak A, Bai R, Slogar E, et al. Evidence for oligodendrocyte progenitor cell heterogeneity in the adult mouse brain. bioRxiv 2020:2020.03.06.981373

45. Binder MD, Xiao J, Kemper D, Ma GZ, Murray SS, Kilpatrick TJ. Gas6 increases myelination by oligodendrocytes and its deficiency delays recovery following cuprizone-induced demyelination. PLoS One 2011;6:e17727

46. Cheng P, Phillips E, Kim SH, Taylor D, Hielscher T, Puccio L, et al. Kinome-wide shRNA screen identifies the receptor tyrosine kinase AXL as a key regulator for mesenchymal glioblastoma stem-like cells. Stem Cell Reports 2015;4:899–913

47. McKinnon RD, Piras G, Ida JA, Jr., Dubois-Dalcq M. A role for TGF-beta in oligodendrocyte differentiation. J Cell Biol 1993;121:1397–407

48. Petersen MA, Ryu JK, Chang KJ, Etxeberria A, Bardehle S, Mendiola AS, et al. Fibrinogen Activates BMP Signaling in Oligodendrocyte Progenitor Cells and Inhibits Remyelination after Vascular Damage. Neuron 2017;96:1003–12 e7

49. Nolan K, Thompson TB. The DAN family: modulators of TGF-beta signaling and beyond. Protein Sci 2014;23:999–1012

50. Pitts RL, Wang S, Jones EA, Symes AJ. Transforming growth factor-beta and ciliary neurotrophic factor synergistically induce vasoactive intestinal peptide gene expression through the cooperation of Smad, STAT, and AP-1 sites. J Biol Chem 2001;276:19966–73

51. Dai ZM, Sun S, Wang C, Huang H, Hu X, Zhang Z, et al. Stage-specific regulation of oligodendrocyte development by Wnt/beta-catenin signaling. J Neurosci 2014;34:8467–73

52. Marques S, van Bruggen D, Vanichkina DP, Floriddia EM, Munguba H, Varemo L, et al. Transcriptional Convergence of Oligodendrocyte Lineage Progenitors during Development. Dev Cell 2018;46:504–17 e7

53. Lykke-Andersen J, Shu MD, Steitz JA. Human Upf proteins target an mRNA for nonsense-mediated decay when bound downstream of a termination codon. Cell 2000;103:1121–31

54. Liu Y, Masson JY, Shah R, O’Regan P, West SC. RAD51C is required for Holliday junction processing in mammalian cells. Science 2004;303:243–6

55. Fekairi S, Scaglione S, Chahwan C, Taylor ER, Tissier A, Coulon S, et al. Human SLX4 is a Holliday junction resolvase subunit that binds multiple DNA repair/recombination endonucleases. Cell 2009;138:78–89

56. Yang P, Kollmeyer TM, Buckner K, Bamlet W, Ballman KV, Jenkins RB. Polymorphisms in GLTSCR1 and ERCC2 are associated with the development of oligodendrogliomas. Cancer 2005;103:2363–72

57. Janke R, Kong J, Braberg H, Cantin G, Yates JR, 3rd, Krogan NJ, et al. Nonsense-mediated decay regulates key components of homologous recombination. Nucleic Acids Res 2016;44:5218–30

58. Marques C, Unterkircher T, Kroon P, Izzo A, Gargiulo G, Kling E, et al. NF1 regulates mesenchymal glioblastoma plasticity and aggressiveness through the AP-1 transcription factor FOSL1. bioRxiv 2019:834531

59. Matos DA, Zhang JM, Ouyang J, Nguyen HD, Genois MM, Zou L. ATR Protects the Genome against R Loops through a MUS81-Triggered Feedback Loop. Mol Cell 2020;77:514–27 e4

60. Wang Y, Waldron RT, Dhaka A, Patel A, Riley MM, Rozengurt E, et al. The RAS effector RIN1 directly competes with RAF and is regulated by 14-3-3 proteins. Mol Cell Biol 2002;22:916–26

61. Sun T, Warrington NM, Luo J, Brooks MD, Dahiya S, Snyder SC, et al. Sexually dimorphic RB inactivation underlies mesenchymal glioblastoma prevalence in males. J Clin Invest 2014;124:4123–33

62. Sugiarto S, Persson AI, Munoz EG, Waldhuber M, Lamagna C, Andor N, et al. Asymmetry-defective oligodendrocyte progenitors are glioma precursors. Cancer Cell 2011;20:328–40

63. Chen G, Kong J, Tucker-Burden C, Anand M, Rong Y, Rahman F, et al. Human Brat ortholog TRIM3 is a tumor suppressor that regulates asymmetric cell division in glioblastoma. Cancer Res 2014;74:4536–48

64. Lein ES, Hawrylycz MJ, Ao N, Ayres M, Bensinger A, Bernard A, et al. Genome-wide atlas of gene expression in the adult mouse brain. Nature 2007;445:168–76

