## Supplementary Figures and Tables for "Premalignant *Nf1, Trp53*-null Oligodendrocyte Precursor Cells Become Stalled in a Heterogeneous State of Replication Stress Before Gliomagenesis"

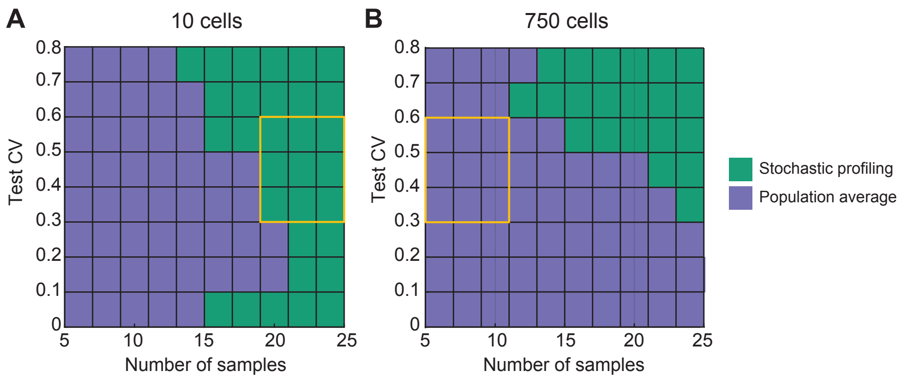


**Supplementary Figure S1.** Bulk analysis and stochastic profiling occupy different experimental regimes. **A** and **B,** Monte-Carlo simulations of stochastic-profiling experiments were simulated with 10-cell pools (**A**) or 750-cell pools (**B**) with the following parameters: $D$ = 5, $F$ = 0.5, $\sigma_{b1}$ = $\sigma_{b2}$ = 0.3. Yellow boxes highlight the range of experimental samples collected for each regime. See (1) for details on the simulation parameters.

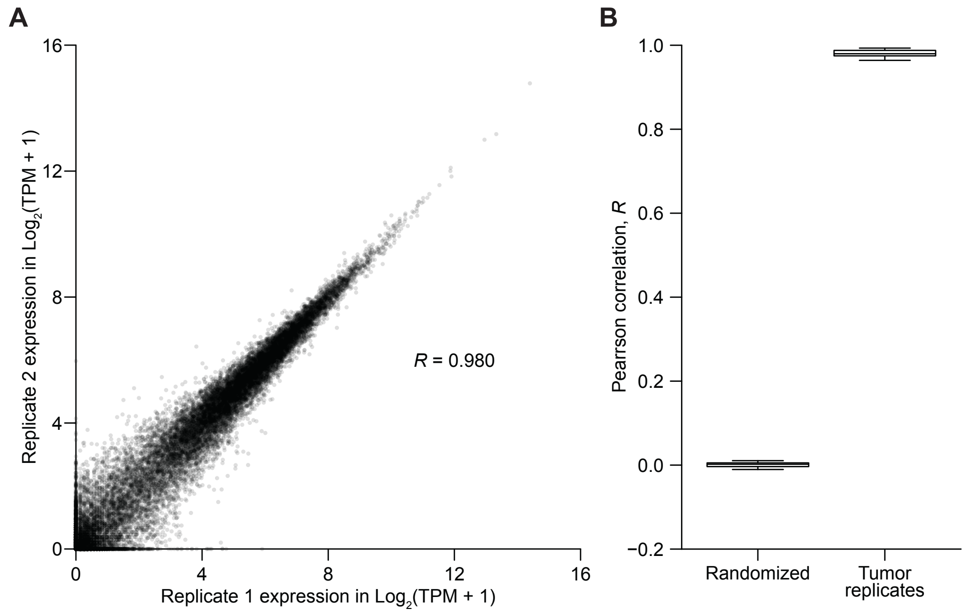


**Supplementary Figure S2.** RNA sequencing of mutant tumors after linear amplification is highly reproducible. **A,** Correlation plot of two 820-cell samples microdissected from the same 150 dpi mutant tumor and amplified independently. **B,** Boxplot of pairwise correlations within 150 dpi mutant tumors. Randomized transcripts are shown as a negative control. Data are from *n* = 7 pairs of independent samples.

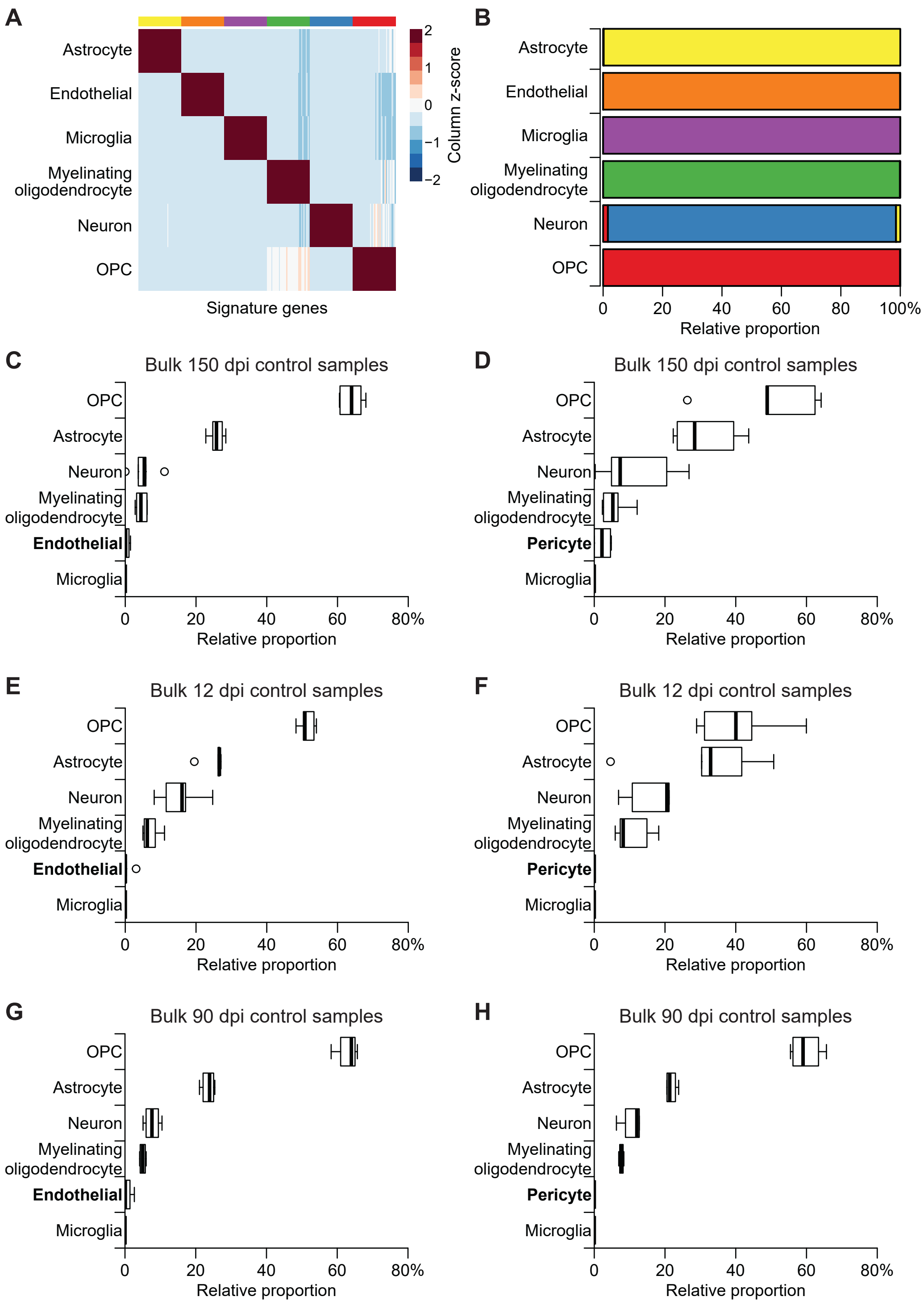
A
Astrocyte
Endothelial
Microglia
Myelinating oligodendrocyte
Neuron
OPC
Signature genes
Column z-score
2
1
0
−1
−2
B
Astrocyte
Endothelial
Microglia
Myelinating oligodendrocyte
Neuron
OPC
0 20 40 60 80 100%
Relative proportion
C
Bulk 150 dpi control samples
OPC
Astrocyte
Neuron
Myelinating oligodendrocyte
Endothelial
Microglia
0 20 40 60 80%
Relative proportion
D
Bulk 150 dpi control samples
OPC
Astrocyte
Neuron
Myelinating oligodendrocyte
Pericyte
Microglia
0 20 40 60 80%
Relative proportion
E
Bulk 12 dpi control samples
OPC
Astrocyte
Neuron
Myelinating oligodendrocyte
Endothelial
Microglia
0 20 40 60 80%
Relative proportion
F
Bulk 12 dpi control samples
OPC
Astrocyte
Neuron
Myelinating oligodendrocyte
Pericyte
Microglia
0 20 40 60 80%
Relative proportion
G
Bulk 90 dpi control samples
OPC
Astrocyte
Neuron
Myelinating oligodendrocyte
Endothelial
Microglia
0 20 40 60 80%
Relative proportion
H
Bulk 90 dpi control samples
OPC
Astrocyte
Neuron
Myelinating oligodendrocyte
Pericyte
Microglia
0 20 40 60 80%
Relative proportion

**Supplementary Figure S3.** OPC enrichment of tdTomato-directed LCM is confirmed by CIBERSORT. **A,** CIBERSORT signature matrix (2) derived from cell type-enriched markers obtained by RNA sequencing (3) and used in **Fig. 2C**. Signature genes were column standardized and hierarchically clustered. **B,** Validation of the derived signature matrix for cleanly deconvolving transcriptomes from sorted populations of the indicated cell types into astrocytes (yellow), endothelial cells (orange), microglia (purple), myelinating oligodendrocytes (green), neurons (blue), and OPCs (red). **C–H,** Relative CIBERSORT deconvolution of normal 150 dpi (**C** and **D**), 12 dpi (**E** and **F**), or 90 dpi (**G** and **H**) control samples with a separate signature matrix using additional RNA sequencing-based markers of pericytes (4) and optimized to swap endothelial cells (**C**, **E**, **G**) for pericytes (**D**, **F**, **H**).

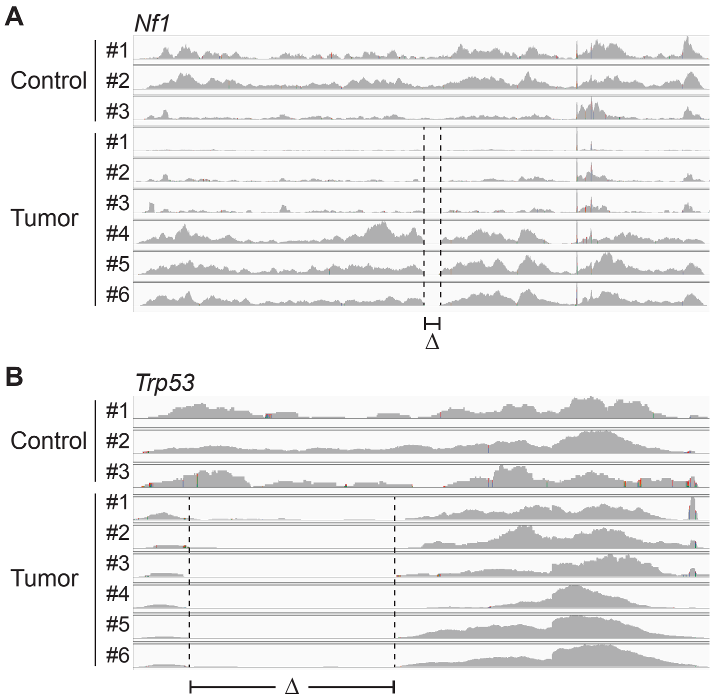


**Supplementary Figure S4.** Differential transcript abundance neglects the functional loss of floxed alleles. **A** and **B,** Deletion of *Nf1* (**A**) and *Trp53* (**B**) exons in mutant tumors confirmed by bulk RNA-seq. The deleted exons (Δ) in the mutant tumors are highlighted. RNA-seq alignment tracks are from *n* = 3 control 150 dpi samples and *n* = 6 mutant tumors.

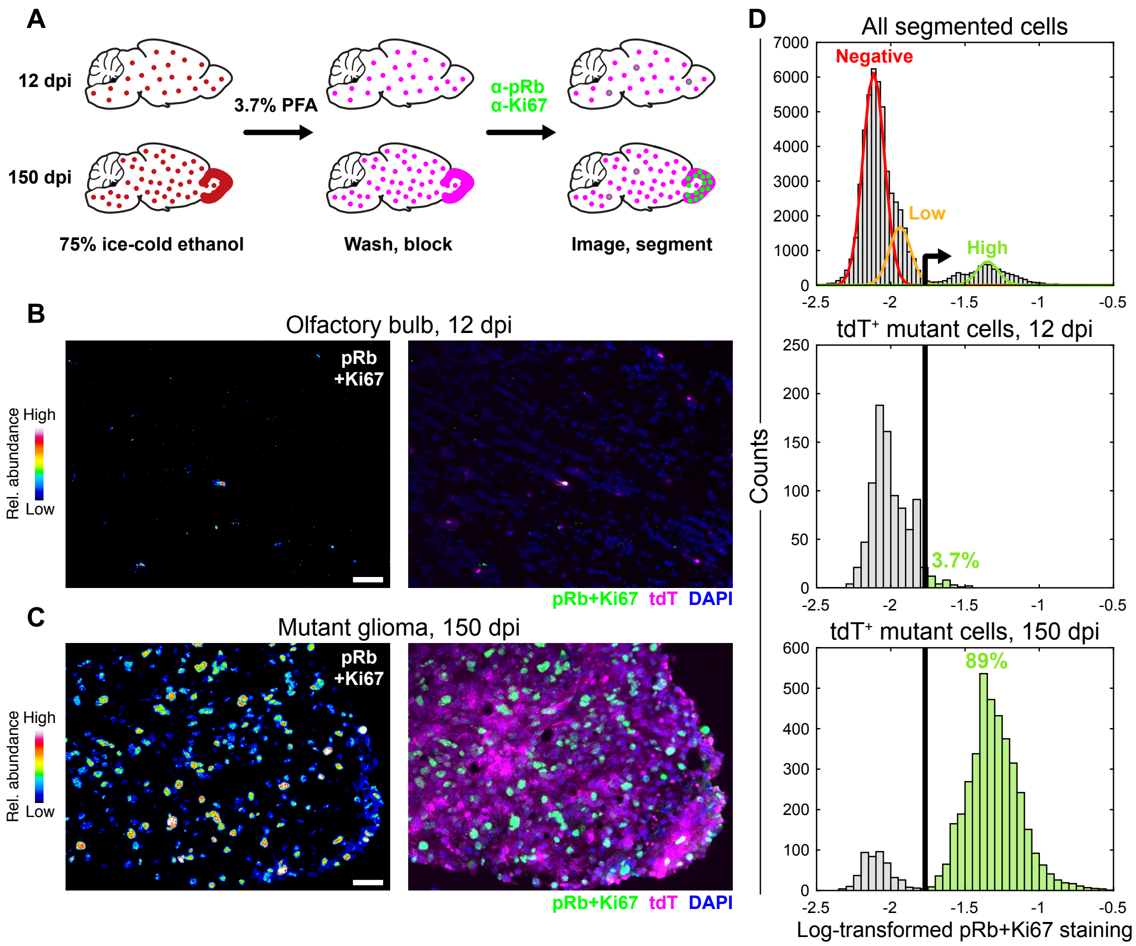


**Supplementary Figure S5.** Mutant cells are not actively proliferating at 12 dpi. **A,** Immunocytochemical procedure for antibody staining that preserves localized tdTomato fluorescence. To monitor proliferation, cells were immunostained for phosphorylated Rb (pRb) and Ki67 in the same fluorescence channel. **B** and **C,** Proliferation is extensive in mutant gliomas at 150 dpi (**C**) but limited among tdTomato$^+$ (tdT$^+$) mutant cells at 12 dpi (**B**). **D,** Summary quantification of pRb+Ki67 immunostaining of 55,479 nuclei (top), 912 tdT$^+$ mutant cells at 12 dpi (middle), or 4359 tdT$^+$ mutant cells at 150 dpi (bottom) from *n* = 4 animals at each time point. The pRb+Ki67 positive gate was set at the 99$^{th}$ percentile of the low subpopulation (orange).

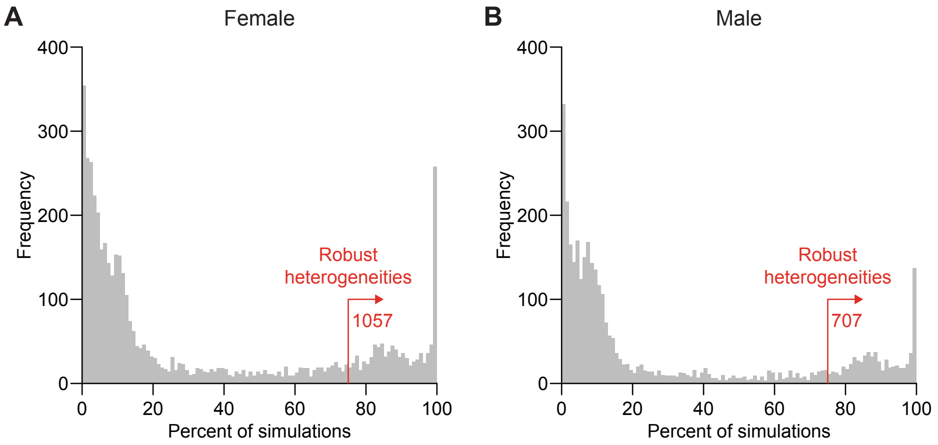


**Supplementary Figure S6.** Leave-one-out cross-validation identifies robust transcriptional heterogeneities in 12 dpi 10-cell samples. **A** and **B,** Cross-validated dispersion analysis of 24 10-cell observations in females (**A**) or males (**B**). Cross-validation runs removed one 10-cell observation and one pool-and-split control, substituting each with replacement from the remaining samples. Each run was analyzed for overdispersion as described (5), and candidates appearing in >75% of cross-validation runs were considered robust heterogeneities.

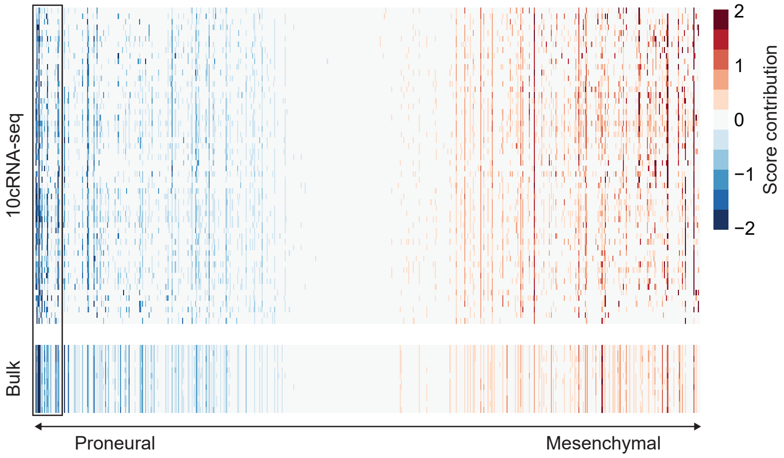


**Supplementary Figure S7.** Contribution of transcripts to the score along the proneural–mesenchymal principal component in 10cRNA-seq samples and bulk samples at 12 dpi. High-abundance transcripts strongly contributing to the proneural score of bulk samples are boxed.

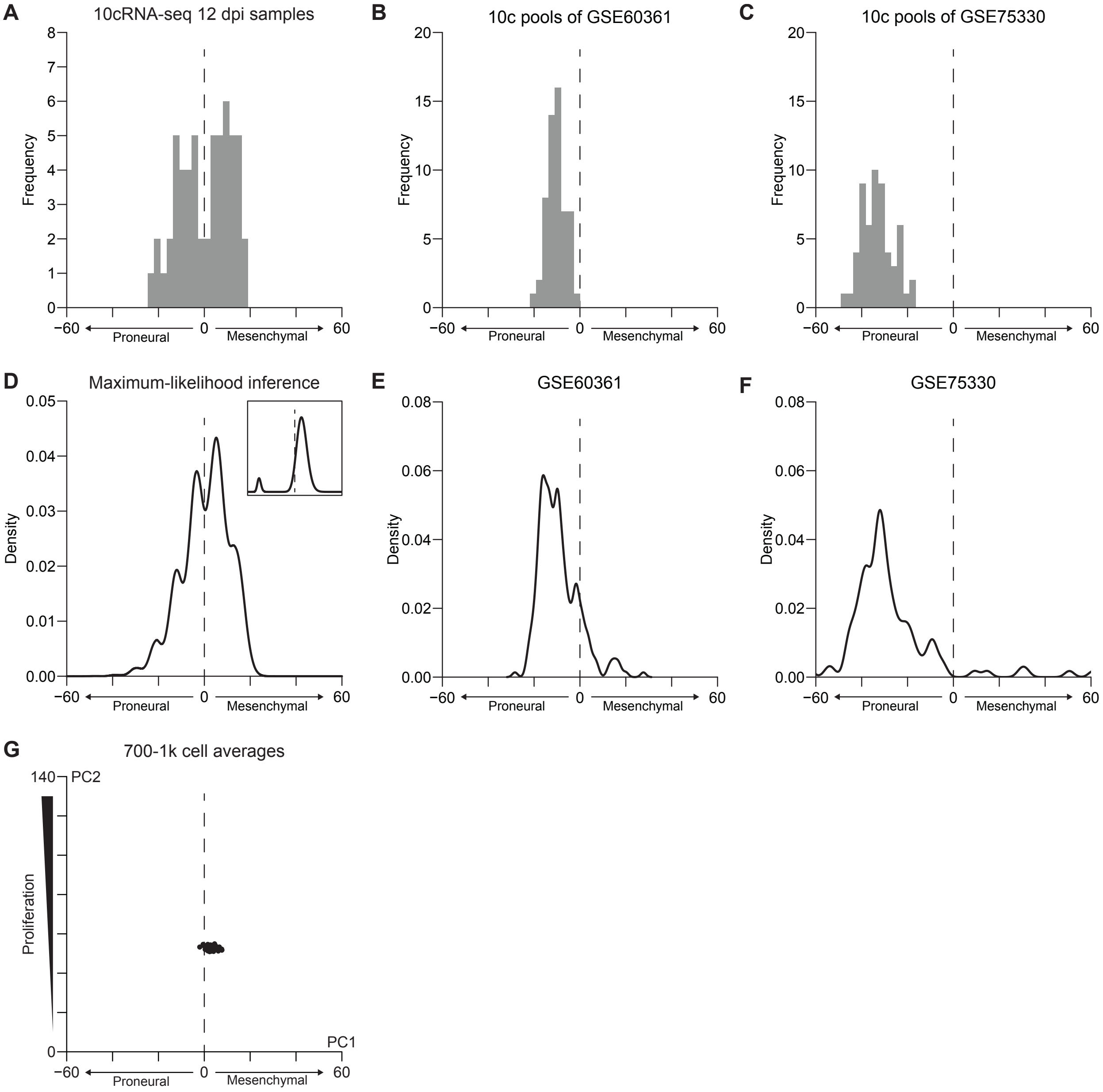


**Supplementary Figure S8.** Deconvolution of single-cell projections from 10-cell data compared to single cells and 10-cell pools of normal OPCs. **A,** Histogram of 12 dpi 10cRNA-seq projections collapsed along the proneural-mesenchymal axis of **Fig. 3C**. **B** and **C,** Simulated 10-cell pools of normal OPCs from two public datasets: GSE60361 (**B**) and GSE75330 (**C**). **D,** Single-cell parameterization of 10-cell probability densities estimated by maximum-likelihood inference (6) based on the 10-cell averages in **A**. The inset shows the proportion and variance of the two underlying single-cell distributions (to be compared with **E** and **F**). **E** and **F,** scRNA-seq distributions of normal OPCs from two public datasets: GSE60361 (**E**) and GSE75330 (**F**). **G,** Simulated projection of 700–1000 cells averaged from the 10-cell projections observed in **Fig. 3C** (see Materials and Methods).

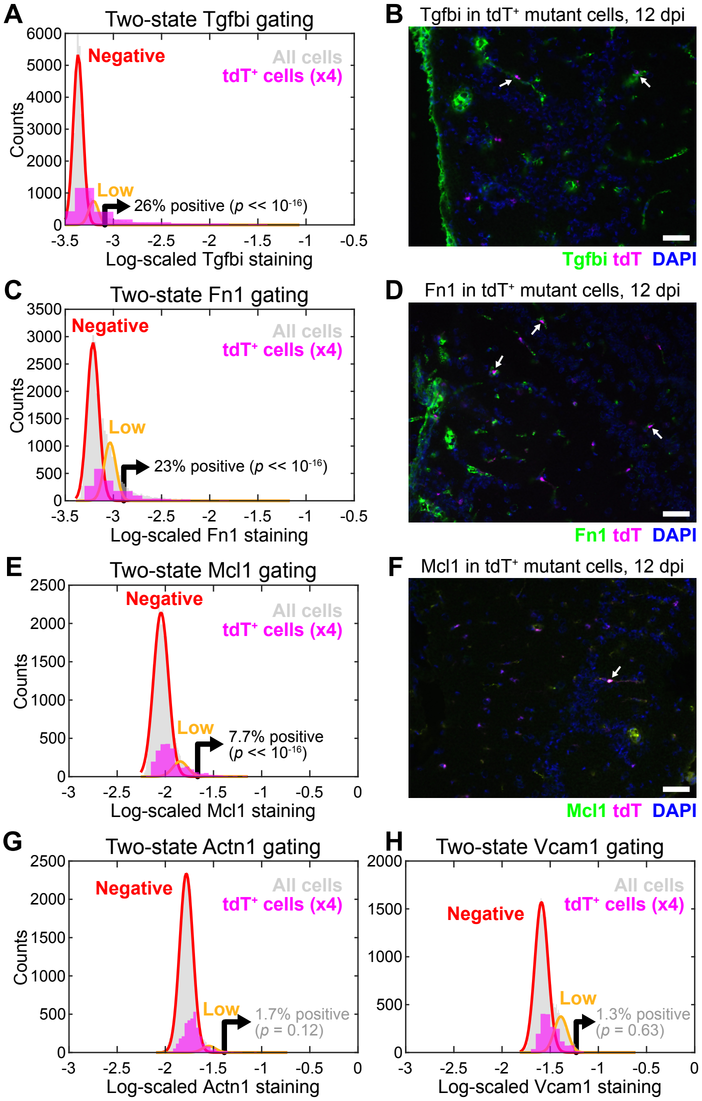


**Supplementary Figure S9.** Mutant cells sporadically express mesenchymal markers at 12 dpi. **A–F,** Quantitative immunocytochemistry (**A**, **C**, and **E**) and representative images (**B**, **D**, and **F**) of 12 dpi cryosections stained for Tgfbi (**A** and **B**), Fn1 (**C** and **D**), or Mcl1 (**E** and **F**). Summary quantifications are from 45,370 (**A** and **E**) or 33,568 (**C**) nuclei (gray) and 627 (**A** and **E**) or 461 (**C**) tdTomato$^+$ (tdT$^+$) mutant cells (magenta) from $n$ = 4 animals. White arrows in **B**, **D**, and **F** indicate pericellular (**B** and **D**) or intracellular (**F**) colocalization. **G** and **H,** Actn1 and Vcam1 are not consistently detected in mutant cells at 12 dpi. Quantitative immunocytochemistry of 12 dpi cryosections stained for Actn1 (**G**) or Vcam1 (**H**). Summary quantifications are from 42,465 (**G**) or 33,568 (**H**) nuclei (gray) and 842 (**G**) or 461 (**H**) tdT$^+$ mutant cells (magenta) from $n$ = 4 animals. In **A**, **C**, **E**, and **G–H**, positive gates were set at the 99$^{th}$ percentile of the low subpopulation (orange), and histogram counts of tdT$^+$ cells were multiplied by four to visualize the distributions more clearly. Significance of the positive subpopulation was assessed by binomial test with a 1% background probability.

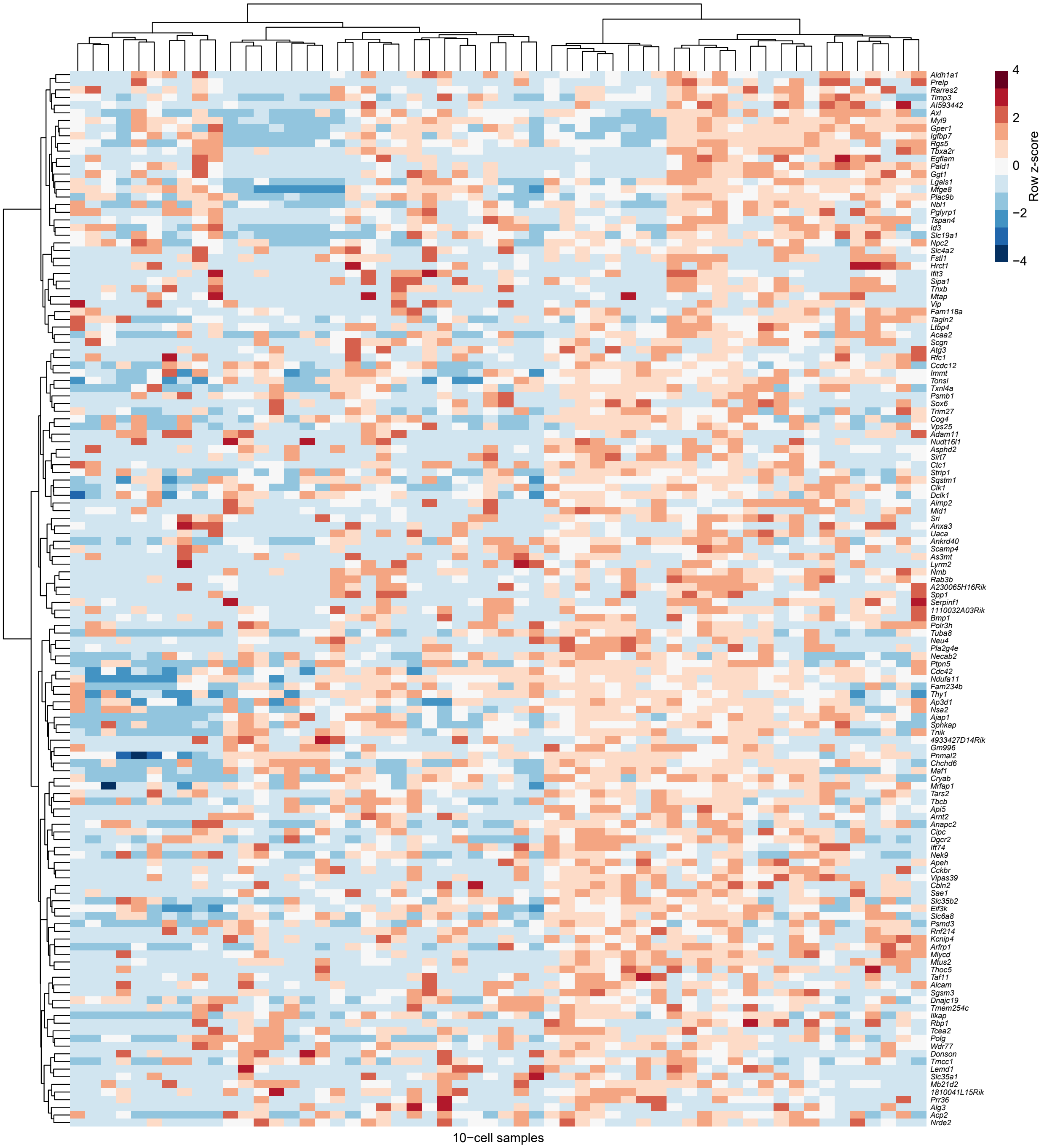


**Supplementary Figure S10.** Sample-by-sample clustergram of RHEGs in mutant OPCs at 12 dpi.

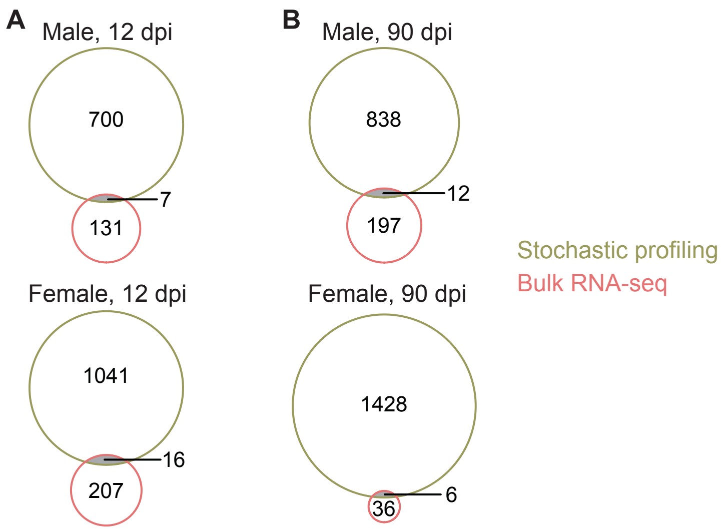


**Supplementary Fig. S11.** Bulk differential expression and single-cell regulatory heterogeneities are not significantly overlapping when separated by sex. **A** and **B,** Venn diagram intersecting candidate heterogeneities and differentially expressed genes for males (upper) and females (lower) at 12 dpi (**A**) and 90 dpi (**B**). No overlaps were statistically significant ($p \geq 0.35$)

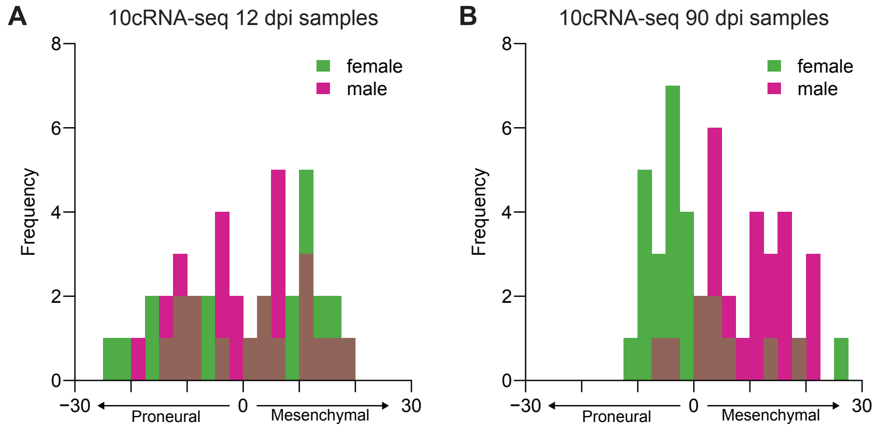


**Supplementary Figure S12.** Histogram of 10cRNA-seq projections along the proneural–mesenchymal axis separated by male and female samples. **A** and **B,** Data from **Fig. 3C** (**A**) and **Fig. 4C** (**B**) were collapsed along PC1 and separated by sex. Histograms summarize 10cRNA-seq projections from *n* = 28 10-cell samples from males (magenta) and 28 10-cell samples from females (green).

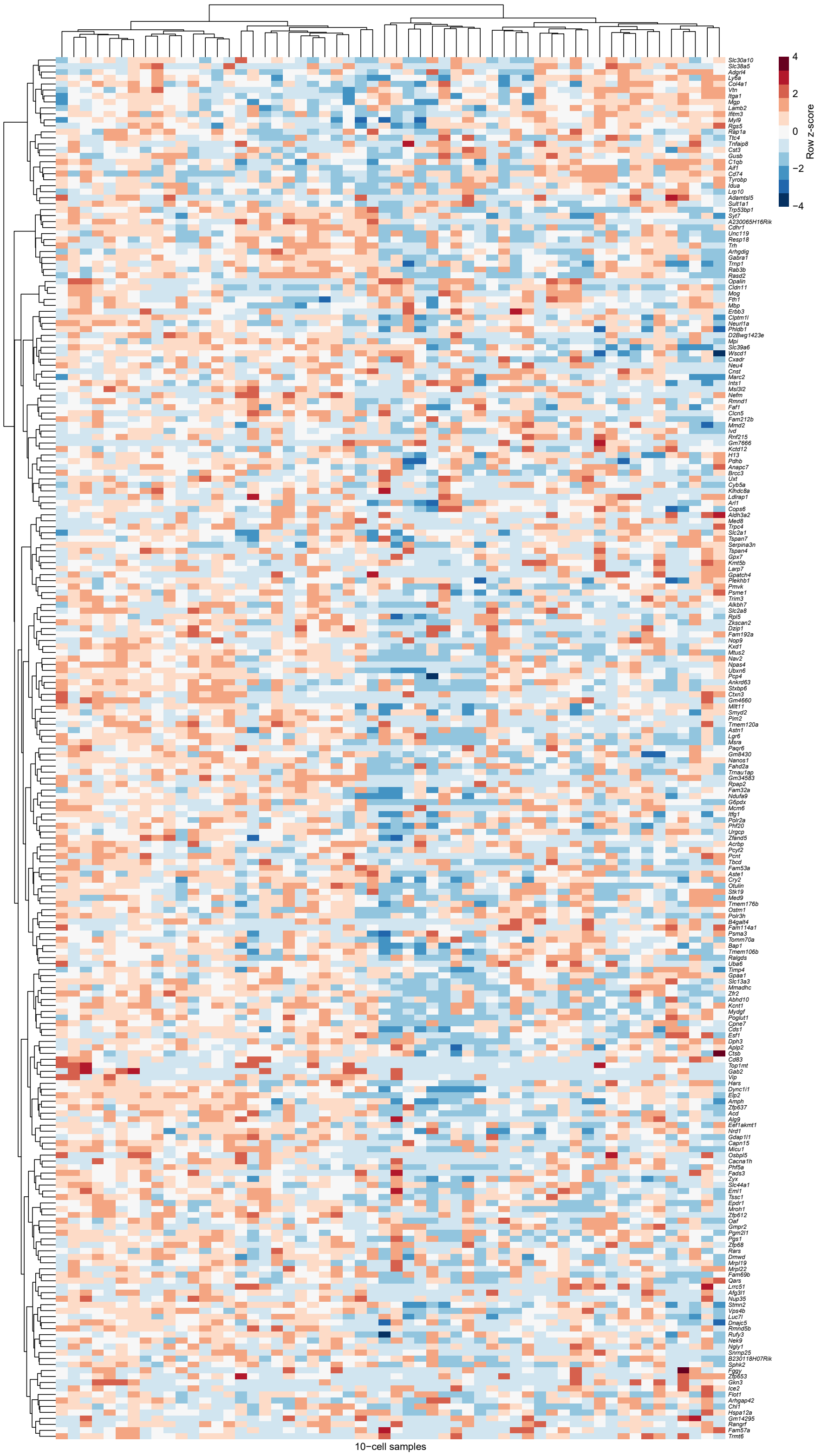


**Supplementary Figure S13.** Sample-by-sample clustergram of RHEGs in mutant OPCs at 90 dpi.

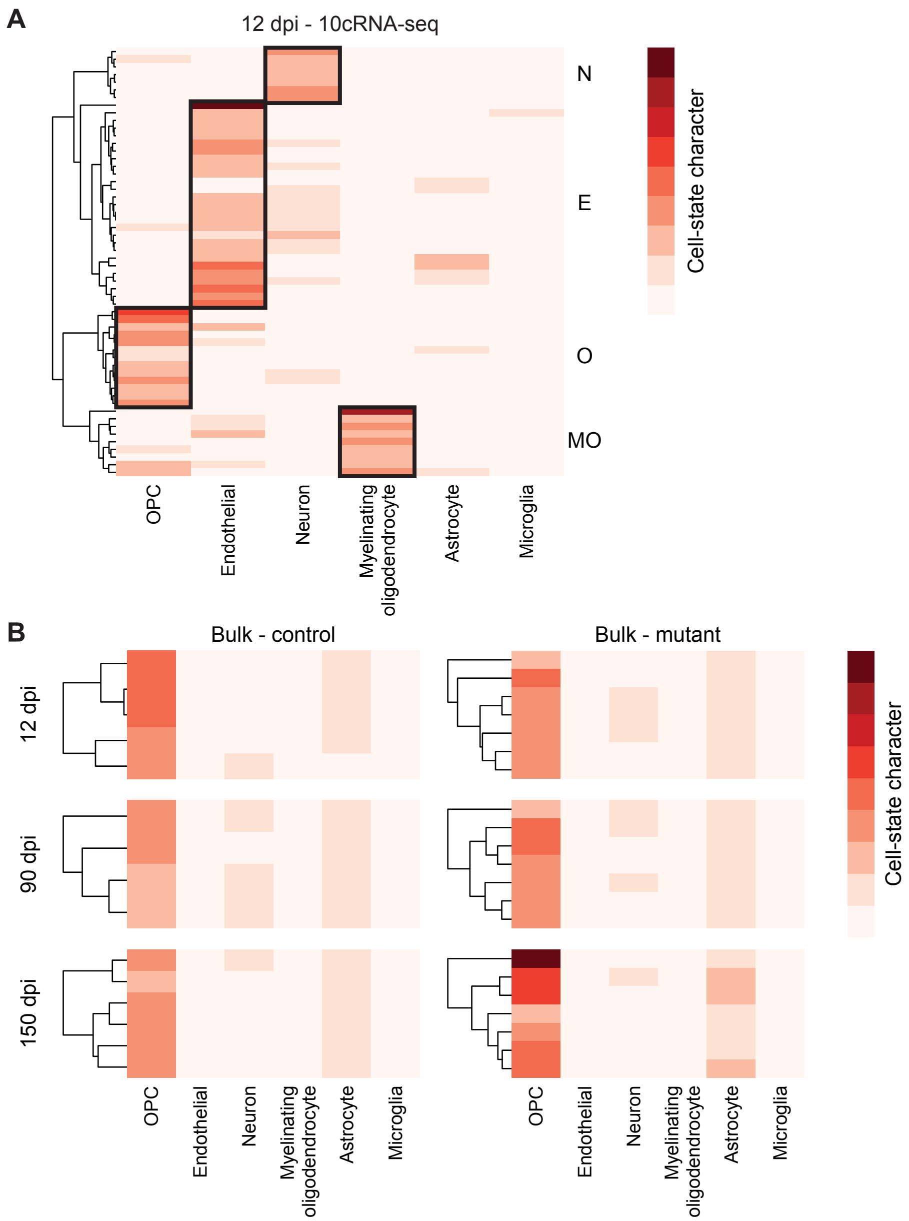


**Supplementary Figure S14.** Absolute CIBERSORT deconvolution of 10cRNA-seq and bulk RNA-seq data. **A,** Unmixing of 10-cell samples at 12 dpi yields defined lineages. Sample groups with substantial cell-state character of OPCs (O), endothelial cells (E), neurons (N), or myelinating oligodendrocytes (MO) are marked. **B,** Absolute unmixing of bulk samples reinforces the predominant O state shown as relative proportions in Supplementary Fig. S3.

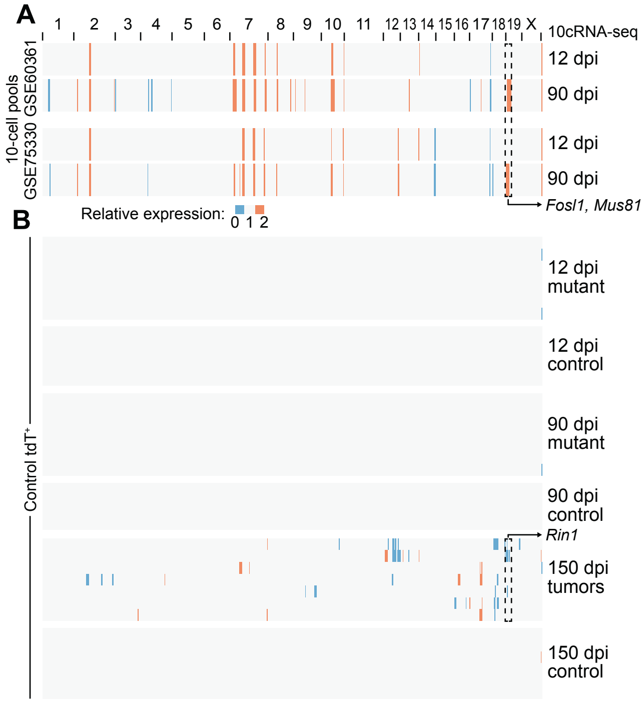


**Supplementary Figure S15.** Stability of inferCNV results. **A,** Localized transcriptional changes inferred using 10-cell pools of GSE60361 (7) or GSE75330 (8) as reference transcriptomes. **B,** Lack of copy number variation detected in mutant bulk samples at 12 dpi and 90 dpi or in control bulk samples at 12 dpi, 90 dpi, or 150 dpi. Data on 150 dpi tumors are reprinted from **Fig. 5K** for comparison.

**Supplementary Table ST1.** List of qPCR primers used in this study.

| Transcript | Forward primer | Reverse primer |
|---|---|---|
| *Gapdh* | GGCATTGCTCTCAATGACAA | GCCTCTCTTGCTCAGTGTCC |
| *Rplp* | ATCTACTCCGCCCTCATCCT | ATGAGGCTCCCAATGTTGAC |
| *Apoe* | GGTTCGAGCCAATAGTGGAA | GCCAGAGAGGTGCTTGAGAC |
| *Cspg4* | CTATCCGTGGCAGCTTCACT | TTCCCTGACGTTCCCTGTAG |
| *Pdgfrb* | GCTGACGGGATTACCACCTC | GAGATGCCAGCTAGCCAACC |
| *Des* | ACACCCAGCCACTTTTCTCC | CGATGACTTGAGCTGGGTTCT |
| *Anpep* | CATGAGGGTGCAAGACCTGG | TTACTATCTCAGCAGCCTGTGC |

**Supplementary File S1.** Hallmark MSigDB enrichments for differentially expressed transcripts between control and mutant bulk samples at 150 dpi (Sheets 1, 2, 7, and 8), 12 dpi (Sheets 3, 4, 9, and 10), and 90 dpi (Sheets 5, 6, 11, and 12). Increased (up, Sheets 1, 3, 5, 7, 9, and 11) and decreased (down, Sheets 2, 4, 6, 8, 10, 12) transcripts were analyzed separately without any fold-change cutoff (Sheets 1–6) or with a two-fold cutoff (fc2.0, Sheets 7–12).

**Supplementary File S2.** Relative transcript abundance changes ($\log_2$ fold change between mutant and control) in bulk samples at 150 dpi (Sheet 1), 12 dpi (Sheet 2), and 90 dpi (Sheet 3). Comparisons are separately broken down according to sex at each dpi time point (Sheets 4–9) .

**Supplementary File S3.** Candidate male- and female-specific regulatory heterogeneities in mutant cells at 12 dpi (Sheet 1) and 90 dpi (Sheet 2). Shared candidates are shown as RHEGs for each day.

### SUPPLEMENTARY REFERENCES


1. Wang L, Janes KA. Stochastic profiling of transcriptional regulatory heterogeneities in tissues, tumors and cultured cells. Nat Protoc **2013**;8:282-301
2. Newman AM, Liu CL, Green MR, Gentles AJ, Feng W, Xu Y*, et al.* Robust enumeration of cell subsets from tissue expression profiles. Nat Methods **2015**;12:453-7
3. Zhang Y, Chen K, Sloan SA, Bennett ML, Scholze AR, O'Keeffe S*, et al.* An RNA-sequencing transcriptome and splicing database of glia, neurons, and vascular cells of the cerebral cortex. J Neurosci **2014**;34:11929-47
4. He L, Vanlandewijck M, Raschperger E, Andaloussi Mae M, Jung B, Lebouvier T*, et al.* Analysis of the brain mural cell transcriptome. Sci Rep **2016**;6:35108
5. Singh S, Sutcliffe MD, Repich K, Atkins KA, Harvey J, Janes KA. Pan-cancer drivers are recurrent transcriptional regulatory heterogeneities in early-stage luminal breast cancer. Cancer Res **2020**:co-submitted
6. Bajikar SS, Fuchs C, Roller A, Theis FJ, Janes KA. Parameterizing cell-to-cell regulatory heterogeneities via stochastic transcriptional profiles. Proc Natl Acad Sci U S A **2014**;111:E626-35
7. Zeisel A, Munoz-Manchado AB, Codeluppi S, Lonnerberg P, La Manno G, Jureus A*, et al.* Brain structure. Cell types in the mouse cortex and hippocampus revealed by single-cell RNA-seq. Science **2015**;347:1138-42
8. Marques S, Zeisel A, Codeluppi S, van Bruggen D, Mendanha Falcao A, Xiao L*, et al.* Oligodendrocyte heterogeneity in the mouse juvenile and adult central nervous system. Science **2016**;352:1326-9
